# The 20S proteasome activator PA28γ controls the compaction of chromatin

**DOI:** 10.1101/716332

**Authors:** Didier Fesquet, David Llères, Charlotte Grimaud, Cristina Viganò, Francisca Méchali, Séverine Boulon, Olivier Coux, Catherine Bonne-Andrea, Véronique Baldin

## Abstract

PA28γ, a nuclear activator of the 20S proteasome, is involved in the degradation of several proteins regulating cell growth and proliferation and in the dynamics of various nuclear bodies, but its precise cellular functions remain unclear. Here, using a quantitative FLIM-FRET based microscopy assay monitoring close proximity between nucleosomes in living human cells, we show that PA28γ controls chromatin compaction. We find that its depletion induces a decompaction of pericentromeric heterochromatin, similarly to that observed upon the knockdown of HP1β, a key factor in heterochromatin structure. We show that PA28γ is present at HP1β-containing repetitive-DNA sequences abundant in heterochromatin and importantly, that HP1β on its own is unable to drive chromatin compaction without the presence of PA28γ. At the molecular level, we show that this novel function of PA28γ is independent of its stable interaction with the 20S proteasome, and most likely depends on its ability to maintain appropriate levels of H3K9me3 and H4K20me3, histone modifications that are both involved in heterochromatin formation. Overall, our results implicate PA28γ as a key factor involved in the higher order structuration of chromatin.

## Introduction

In eukaryotic cells, the differential organization of chromatin into euchromatin and heterochromatin determines genome compaction and activity in the nucleus. Whereas euchromatin is a relaxed state that is generally transcriptionally active, heterochromatin exhibits a dense organizational state throughout interphase, with relatively low transcription levels and an enrichment of repetitive DNA sequences such as satellite repeats, transposable elements and ribosomal DNA (Lippman et al., 2004; Nishibuchi and Nakayama, 2014; Saksouk et al., 2015; Janssen et al., 2018). Heterochromatin is paramount to the stability of eukaryotic genomes. Indeed, loss of control over these repetitive DNA sequences, including mutations produced by the integration or excision of transposable elements and recombination between repeats, can lead to transcriptional perturbation and DNA recombination, all of which events are at the root of oncogenic transformation (Ayarpadikannan and Kim, 2014; Klement and Goodarzi, 2014).

Multiple pieces of evidence from genetic and cell biology studies point to an important involvement of the Heterochromatin Protein-1 (HP1) family of chromodomain proteins (Maison and Almouzni, 2004; Verschure et al., 2005) and trimethylation of histone H3 lysine 9 (H3K9me3) (Martin and Zhang, 2005; Saksouk et al., 2015) and histone H4 lysine 20 (H4K20me3) (Schotta et al., 2004; Oda et al., 2009; Beck et al., 2012; Bosch-Presegue et al., 2017) in establishing and maintaining heterochromatic states. These histone methylation marks serve as molecular anchors for HP1 proteins, notably HP1β, which are required for heterochromatin compaction and silencing (Lachner et al., 2001; Thiru et al., 2004; Dambacher et al., 2013; Bosch-Presegue et al., 2017; Machida et al., 2018). However, the mechanism by which HP1 folds chromatin-containing H3K9me3-H4K20me3 into higher-order structures has not been fully elucidated.

Proteasome-mediated protein degradation is a central pathway that controls the stability and function of numerous proteins in most cellular processes (Collins and Goldberg, 2017). Proteasomes comprise a family of protein complexes resulting from the association of different regulators/activators with the catalytic core, called the 20S proteasome (Rechsteiner and Hill, 2005; Coux et al., 2020). Among their many functions, it is now well established that proteasome complexes are associated with chromatin and enriched at specific sites in the genome (Geng and Tansey, 2012; Kito et al., 2020), thereby suggesting a direct role for chromatin-associated proteasome complexes in genomic processes (McCann and Tansey, 2014).

Among the nuclear 20S proteasome regulators, the homoheptamer PA28γ (also known as REGγ, 11Sγ or Ki antigen) (Ma et al., 1992; Wilk et al., 2000; Mao et al., 2008) promotes the proteasomal degradation of growth-related proteins including the cyclin-dependent kinase inhibitors p21, p19 and p16 and c-Myc (Chen et al., 2007; Li et al., 2007; Li et al., 2015), as well as several important regulatory proteins including steroid receptor coactivator 3 (SRC-3), SirT7 and p53 (Li et al., 2006; Sun et al., 2016; Zhang and Zhang, 2008). Consistent with this, and despite the fact that PA28γ-20S proteasome complexes constitute only a minor fraction (less than 5%) of the whole proteasome population (Fabre et al., 2014), PA28γ is important for cell growth and proliferation. Indeed, PA28γ-knockout mice show a decrease in body size (Murata et al., 1999; Barton et al., 2004) and derived MEF cells display reduced growth and proliferation, increased apoptosis and a slower G1 to S-phase transition. Besides its role in the degradation of growth-related proteins, PA28γ contributes to the control of cell nuclear architecture, since it is involved in the regulation of the dynamics of various nuclear bodies, including Cajal bodies (CBs) (Cioce et al., 2006; Jonik-Nowak et al., 2018), nuclear speckles (NSs) (Baldin et al., 2008), and promyelocytic leukemia bodies (PMLs) (Zannini et al., 2009). PA28γ has also been linked to chromosome stability (Zannini et al., 2008) and DNA repair (Levy-Barda et al., 2011), suggesting a potential role of this proteasome activator in the regulation of chromatin structure.

In this study, we highlight an unsuspected, likely proteasome-independent, function of PA28γ in the control of chromatin compaction. Our investigations reveal that PA28γ is associated with chromatin, notably with repetitive DNA sequences abundant in heterochromatin, and importantly, is required to sustain HP1β-dependent chromatin compaction. Furthermore, we show that PA28γ is necessary to maintain the levels of the H3K9me3 and H4K20me3 heterochromatic marks, thereby establishing PA28γ as an important new regulator of heterochromatin structure.

## Results

### PA28γ controls chromatin compaction in living cells

The involvement of PA28γ in the organization of intra-nuclear structures and the maintenance of chromosome stability suggest that PA28γ could play a key role in the regulation of chromatin structure as well. To explore this hypothesis, we performed quantitative FLIM-FRET (Fluorescence Lifetime Imaging Microscopy-Förster Resonance Energy Transfer) measurements of chromatin compaction at the nanometer-scale in living HeLa cells inactive or not for PA28γ. For this, we established a stable CRISPR/Cas9 PA28γ knockout HeLa cell line (Fig. 1A), expressing either H2B-GFP alone (HeLa^H2B-GFP^-KO-PA28γ) or both H2B-GFP and mCherry-H2B (HeLa^H2B-2FPs^-KO-PA28γ). In these cell lines, PA28γ depletion affected neither H2B-GFP nor mCherry-H2B expression levels, as analyzed by immunoblot and microscopy approaches (Fig. S1A,B). FRET was measured between the fluorophore-tagged histones incorporated into the chromatin: in this assay an increase in FRET efficiency corresponds to an increase in the occurrence of close proximity (<10 nm) between nucleosomes (Lleres et al., 2009). In wild-type (WT) HeLa^H2B-2FPs^ cells, a heterogeneous FRET efficiency map was apparent throughout interphase nuclei on representative images using continuous pseudocolors (Fig. 1B). We found that the areas associated with the highest FRET values (red-orange population) decreased in KO-PA28γ cells (Fig. 1B). This effect was confirmed by the determination of the mean FRET efficiency percentage, which shows a major reduction in the level of chromatin compaction compared to the WT cells (Fig. 1C). As a positive control for chromatin decompaction, we treated HeLa^H2B-2FPs^ cells with Trichostatin A (TSA), an inhibitor of histone deacetylases used to induce large-scale chromatin decompaction (Lleres et al., 2009; Otterstrom et al., 2019). As expected, after 24 hours of TSA treatment, the mean FRET efficiency percentage dropped drastically, consistent with a massively decompacted interphase chromatin (Fig. S1C). By extracting the FRET efficiency distribution curves related to the FRET efficiency map of individual nuclei in both WT and KO-PA28γ cell lines, we found that the loss of PA28γ mainly caused a marked reduction of the high FRET population corresponding to high levels of chromatin compaction (Fig. 1D, black curve w blue curve). In contrast, the low-FRET population corresponding to chromatin regions with the lowest degree of chromatin compaction remained poorly affected.

**Figure 1.**
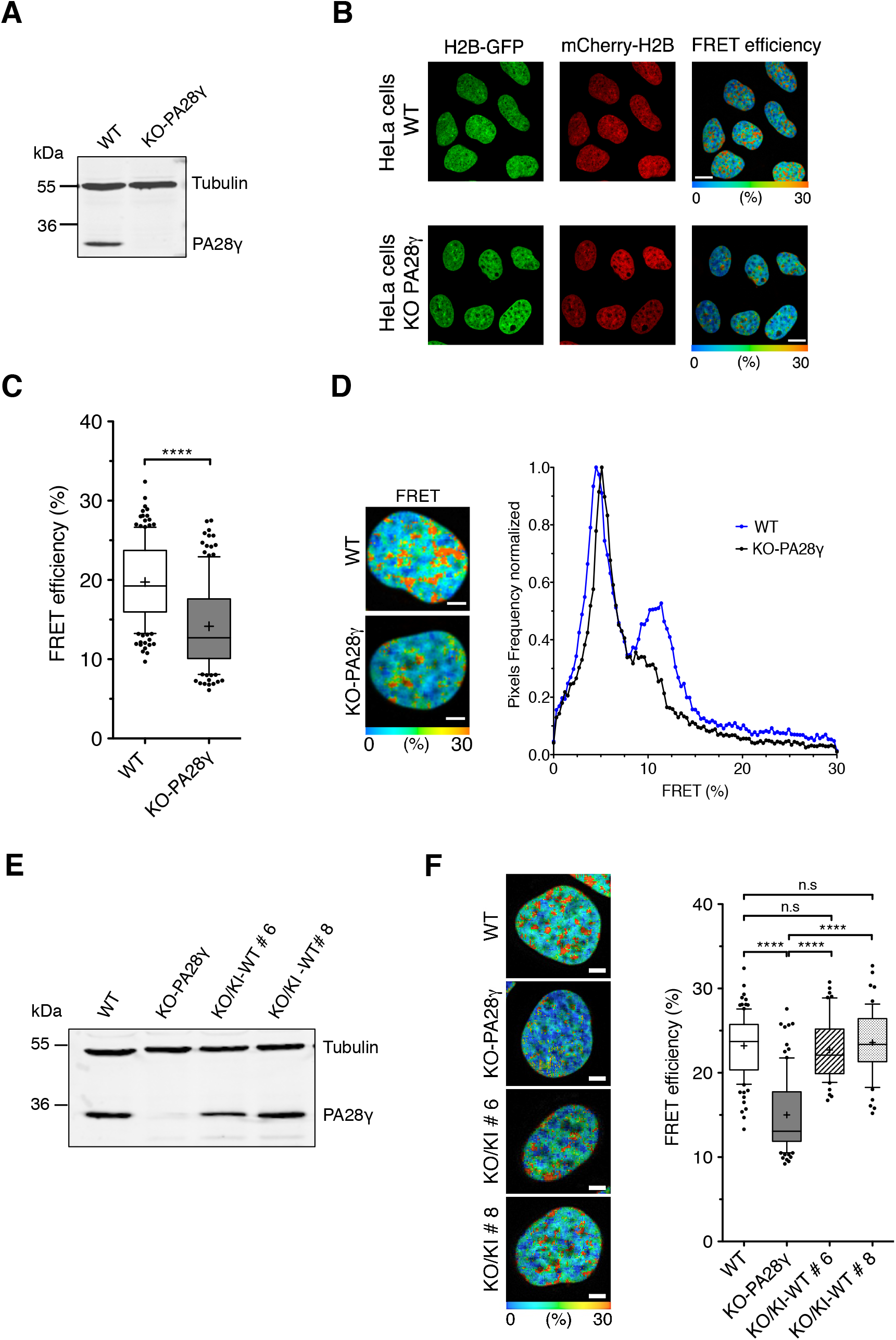
PA28γ controls chromatin compaction. **A.** Immunoblot analysis of PA28γ expression level in total extracts from parental (WT) and PA28γ-knockout (KO-PA28γ) HeLa^H2B-2FPs^ cells. Tubulin was used as a loading control. **B.** FRET analysis in asynchronous interphase parental (WT) and PA28γ-knockout (KO-PA28γ) HeLa^H2B-FPs^ cells. FLIM-FRET measurements were performed and the spatial distribution of the FRET efficiency is represented in a continuous pseudo-color scale ranging from 0 to 30%. Scale bars, 10 μm. **C.** Statistical analysis of the mean FRET efficiency percentage in WT and KO-PA28γ HeLa^H2B-2FPs^ nuclei, presented as box-and-whisker plots. The thick line represents median, the boxes correspond to the mean FRET values above and upper of the median, with the whiskers covering the 10-90 percentile range. Data represent the means ± SD from 4-6 independent experiments, the total number of cells analyzed is *n* = 154 nuclei (WT) and *n* = 132 nuclei (KO-PA28γ), **** *p* < 0.0001 (Student’s *t*-test). **D.** Spatial distribution of the FRET efficiency (percentage) in representative WT and KO-PA28γ HeLa^H2B-2FPs^ nuclei. The FRET percentage distribution is depicted in a continuous pseudo-color scale ranging from 0 to 30% (left panel). Scale bars, 10 μm. FRET distribution graph shows distinct populations of FRET efficiency in WT and KO-PA28γ cells (blue and black curves, respectively) (right panel). **E.** Immunoblot analysis of PA28γ expression level in total extracts from parental (WT), PA28γ-knockout (KO-PA28γ) HeLa^H2B-2FPs^ cells and two independent clones of HeLa^H2B-2FPs^ cells knocked out for PA28γ in which wild-type PA28γ was stably re-expressed (KO/KI-WT #6, KO/KI-WT #8). Tubulin was used as a loading control. **F.** Spatial distribution of the FRET efficiency (percentage) in representative WT, KO-PA28γ and KO/KI-WT #6, KO/KI-WT #8 HeLa^H2B-2FPs^ nuclei. The FRET percentage distribution is depicted as in D. Scale bars, 10 μm. Quantification of the mean FRET efficiency was represented as box-and-whisker plots. Data represent the means ± SD from 3 independent experiments, the total number of cells analyzed is *n* = 102 nuclei (WT), *n* = 90 (KO-PA28γ), *n* = 53 (KO/KI-WT #6), *n* = 54 (KO/KI-WT #8). n.s, not significant, **** *p* <0.0001 (Student’s *t*-test).

To ascertain that the observed chromatin decompaction was due to the absence of PA28γ, we re-expressed PA28γ in PA28γ-KO cell lines (two different clones named KO/KI-WT#6 and #8 were selected) at a level comparable to that of the endogenous protein (Fig. 1E). Remarkably, for KO/KI-WT#6 and #8 clones, the FRET efficiency was restored to values similar to WT cells (Fig. 1F) indicating the re-establishment of normal chromatin compaction. Thus, these results show that PA28γ plays an important role in regulating the compaction of chromatin in interphase cells, with a particular impact on the most condensed chromatin regions.

### PA28γ controls chromatin compaction independently from its interaction with the 20S proteasome

Given that PA28γ has functions that are either proteasome binding-dependent (Li et al., 2007; Levy-Barda et al., 2011) or proteasome binding-independent (Zannini et al., 2008; Zhang and Zhang, 2008), we asked whether the role of PA28γ in the regulation of chromatin compaction requires its interaction with the 20S proteasome. For this purpose, a mutant of PA28γ deleted of its C-terminal 14 amino acids (named ΔC), which is unable to bind and to activate the 20S proteasome (Ma et al., 1993; Förster et al., 2005; Zhang and Zhang, 2008; Zannini et al., 2008), was stably expressed at a physiological level in HeLa^H2B-2FPs^-KO-PA28γ cells (named KO/KI-ΔC) (Fig. 2A). The inability of this PA28γ mutant to bind the 20S proteasome was confirmed by co-immunoprecipitation experiments from cell extracts treated or not with the proteasome inhibitor MG132, known to increase the association between PA28γ and the 20S proteasome (Welk et al., 2016). As shown in Fig. 2B and Fig. S2, the 20S proteasome was detected by the presence of its α4 subunit in PA28γ immunoprecipitations from HeLa^H2B-2FP^ WT and KO/KI-WT cell extracts, but not in KO/KI-ΔC and KO-PA28γ cells. Chromatin compaction was then analyzed by FLIM-FRET in living asynchronous cells. We found that expression of the PA28γ-ΔC mutant restored the level of chromatin compaction in PA28γ-KO cells to a FRET efficiency value (24.7%) similar to that observed in WT cells (23.03%) (Fig. 2C). These results demonstrate that compaction of chromatin requires PA28γ, but not its stable interaction with the 20S proteasome.

**Figure 2.**
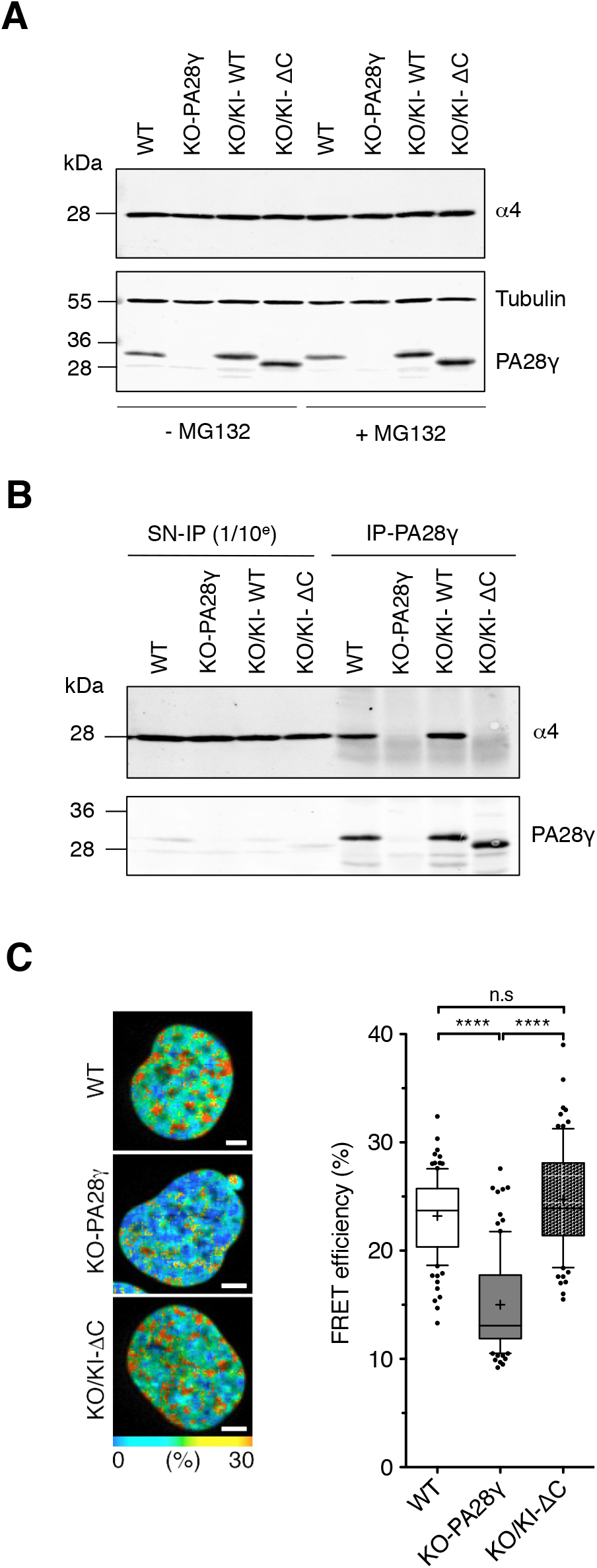
PA28γ interaction with the 20S proteasome is not required for chromatin compaction. **A.** Parental HeLa^H2B-2FPs^ (WT), PA28γ-knockout (KO-PA28γ) cells and KO cells re-expressing the wild-type (KO/KI-WT#8) form or the ΔC-mutant (KO/KI-ΔC) of PA28γ were treated, or not, for 2 hours with MG132 (25 μM), and whole-cell extracts were analyzed by immunoblot using the antibodies indicated. **B.** Cell extracts from cell lines treated with MG132 as in A were subjected to immunoprecipitation using anti-PA28γ antibodies. Immunoblots of the supernatant (SN-IP) and the pull-down (IP-PA28γ) from whole-cell extracts were probed with the antibodies indicated. **C.** Spatial distribution of the FRET efficiency (percentage) in representative WT, KO-PA28γ and KO cells re-expressing the ΔC-mutant (KO/KI-ΔC) HeLa^H2B-2FPs^ nuclei. The FRET percentage distribution is depicted as in Fig. 1C (left panel). Scale bars, 10 μm. Quantification of the FLIM-FRET measurements. Data represent the means ± SD from 3 independent experiments, the total number of cells analyzed is *n* = 102 nuclei (WT), *n* = 90 nuclei (KO-PA28γ), *n* = 83 nuclei (KO/KI-ΔC). n.s, not significant, **** *p* < 0.0001, (Student’s *t*-test).

### PA28γ controls pericentromeric heterochromatin compaction

Since the loss of PA28γ mainly affects chromatin regions with a high-FRET efficiency (Fig. 1D), we hypothesized that PA28γ might play a role in the regulation of highly-compacted heterochromatin. To explore further this possibility, we used a previously-described U2OS cell clone (F42B8) carrying *lacO* DNA repeats stably integrated within constitutive heterochromatin, at a pericentromeric region described as one of the most compacted chromatin domains in the nucleus (Jegou et al., 2009). This *lacO* array forms a single heterochromatic locus that can be visualized in cells following the transient expression of the GFP-LacI construct. The GFP signal allows us to measure the area occupied by the *lacO* locus and thus to quantify the variations of its accessibility and compaction state. As a control of measurable heterochromatin decompaction, we first examined the effect of the depletion of a known regulator of heterochromatin, HP1β (Maison and Almouzni, 2004; Bosch-Presegue et al., 2017), on *lacO* array compaction. These cells were transfected with siRNAs directed against HP1β (si-HP1β) or luciferase (si-Luc), and with a GFP-LacI-expressing construct. The efficiency of si-HP1β was verified by immunoblot (Fig. 3A, upper panel), and changes in heterochromatin compaction state were monitored 48 hours posttransfection by fluorescence microscopy (Fig. 3A, lower panel). *LacO* locus appeared as a small dot with a surface area that was not significantly affected by the transfection of si-Luc (0.390 ± 0.045 μm^2^ *vs* 0.370 ± 0.052 μm^2^ in control cells). Upon HP1β knockdown, we observed a significant increase of the GFP-LacI dot surface area (0.730 ± 0.069 μm^2^). This corresponds to an expansion of the surface area occupied by the *lacO* DNA repeats due to heterochromatin decompaction (Fig. 3B). Then, we examined the effect of PA28γ knockdown (Fig. 3C,D). Upon PA28γ-depletion we observed a significant increase in the GFP-LacI dot surface area (0.636 ± 0.014 μm^2^ *vs* 0.370 ± 0.052 μm^2^ and 0.443 ± 0.011 μm^2^ in control cells and si-Luc treated cells, respectively) (Fig. 3D). Thus, as observed in cells depleted for HP1β, the pericentromeric heterochromatin is significantly decompacted in the absence of PA28γ, supporting a role for PA28γ in the control of heterochromatin compaction.

**Figure 3.**
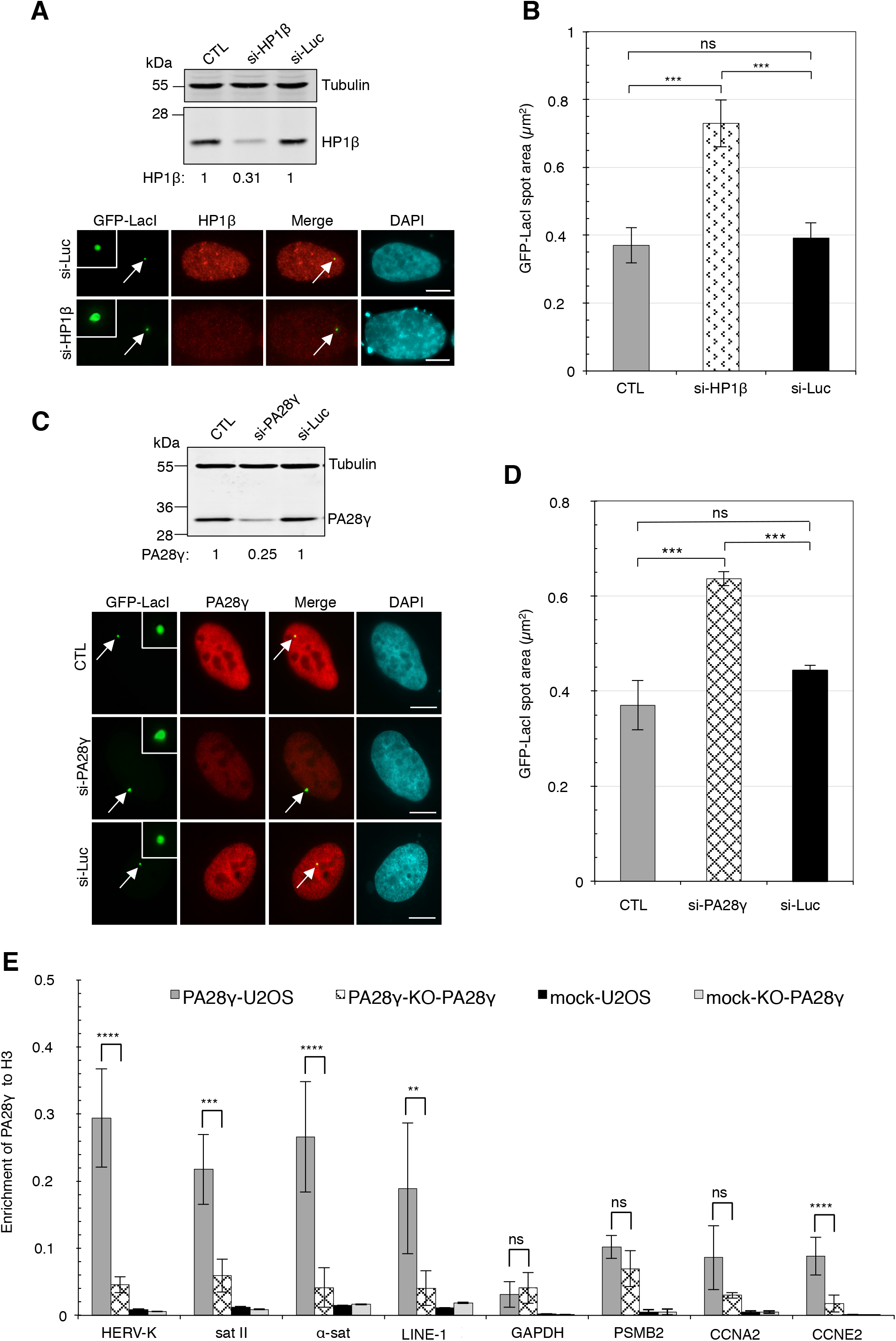
PA28γ depletion induces a decompaction of pericentromeric heterochromatin and PA28γ is present on heterochromatin DNA sequences. **A.** U2OS-LacO cells, treated or not with si-HP1β or si-Luc, were transiently transfected with GFP-LacI construct the same day and were recovered 48 hours later. Proteins were analyzed by immunoblotting. The relative abundance of HP1β in the extracts was quantified using ImageJ software and normalized to tubulin (upper panel). Cells on coverslips were immunostained with anti-HP1β (red) and the GFP signal was imaged in parallel (green). DNA was stained with DAPI (cyan). Representative fluorescence and immunofluorescence images of Z-stack projections of U2OS-LacO cells are shown. Magnified views of GFP-LacI spot are shown in inserts. Scale bars, 10 μm. **B.** Quantitative analysis of the decompaction of the LacO array. Z-stack images were acquired on U2OS-LacO cells treated as in (A), and the area of the GFP-LacI signal was quantified on a Z-projection using ImageJ software (see Materials and Methods). Data represent the means ± SD from three biological repeats, numbers of analyzed nuclei with GFP-LacI spot were *n* = 30, *n* = 28 and *n* = 27 in control cells (CTL), si-HP1β or si-Luc treated cells, respectively. (ns, not significant, *p* = 0.2503 (CTL/si-Luc), *** *p* = 0.0003 and 0.001 for (CTL/ si-HP1β) and (si-HP1β/si-Luc) respectively, *p*-values were determined by a 2-way ANOVA test). **C.** U2OS-LacO cells, treated or not with a si-PA28γ or si-Luc, were transiently transfected with GFP-LacI construct the same day, recovered 48 hours later and cells were analyzed as in A. Immunostaining was performed with antibodies raised against PA28γ (red) and the GFP signal was imaged in parallel (green). DNA was stained with DAPI (cyan). Representative fluorescence and immunofluorescence images of Z-stack projections of U2OS-LacO cells are shown. Magnified views of the GFP-LacI spot are shown in inserts. Scale bars, 10 μm. **D.** Quantitative analysis of the decompaction of the LacO array. Z-stack images were acquired on U2OS-LacO cells treated as in C, and the area of the GFP-LacI signal was quantified as in B. Data represent the means ± SD from three biological repeats, numbers of analyzed nuclei with GFP-LacI spot were *n* = 30, *n* = 31 and *n* = 29 in control cells (CTL), si-PA28γ or si-Luc treated cells, respectively. (ns, not significant, *\*\*\* p* = 0.0002; *\*\*\* p* = 0.00013, values were determined by Tukey’s multiple comparisons test). **E.** ChIP-qPCR analysis of PA28γ levels at different repetitive elements located in heterochromatin or in the promoter of actively transcribed genes (as indicated on the x-axis) in wild-type (WT) *versus* KO-PA28γ U2OS cells (right panel). Data are represented as relative enrichment of PA28γ antibody *versus* histone H3 control, as shown on the y-axis. Data are means ± SEM (*n* = 5). Significance was calculated by Student’s *t*-test, ns = not significant, *(p* = 0.42531, *p* = 0.18602, *p* = 0.2395 for GAPDH, PSMB2 and CCNA2, respectively), *\*\*p* < 0.01 *(p* = 0.0046, LINE-1), *\*\*\*p* < 0.001 *(p* = 0.00011, Sat II), \*\*\*\**p* < 0.0001 *(p =* 2.09E-05, *p* = 5.15E-05 and *p* = 2.08E-07 for HERV-K, α-Sat and CCNE2 respectively).

The effect of PA28γ depletion on heterochromatin compaction prompted us to investigate whether PA28γ might associate with chromatin comprising repetitive sequences characteristic of heterochromatin, such as interspersed (HERV-K), pericentromeric (Satellite II and α Satellite) and Major Satellite (LINE-1) DNA repetitive sequences (Padeken et al., 2015). For this, quantitative chromatin immunoprecipitation (ChIP-qPCR) experiments were performed on parental (WT) *versus* KO-PA28γ U2OS cells, previously characterized (see Fig. S3A and Jonik-Nowak et al., 2018). Since ChIP-qPCR experiments were normalized to histone H3, we first verified that PA28γ depletion did not affect histone H3 expression levels by immunoblot.

As shown in Fig. S3B,C, no variation in expression level of histone H3 was observed in U2OS-KO-PA28γ cells. The same results were obtained for histone H1 and HP1β proteins. ChIP-qPCR experiments revealed that PA28γ was enriched at all four heterochromatin sequences tested (Fig. 3E), as was HP1β (Fig. S3D). By comparison, we analyzed the PA28γ association with sequences located in the promoter of four active transcribed genes *(GAPDH, PSMB2, CCNA2* and *CCNE2).* Of the four euchromatin sequences tested, PA28γ was only detected at the promoter of the cyclin E2 gene, albeit at a much lower level (Fig. 3E), suggesting that its binding is not restricted to heterochromatin regions. Taken together, these results show that PA28γ is a chromatin-binding protein controlling the state of heterochromatin compaction.

### A fraction of PA28γ co-localizes with HP1β

The results described above led us to explore whether PA28γ co-localizes with regulators of heterochromatin establishment such as HP1β. To verify this, we performed immunostaining against endogenous HP1β and PA28γ proteins in U2OS cells. As both are very abundant nuclear proteins and PA28γ displays a diffuse nuclear distribution as well (Masson et al., 2003; Wojcik et al., 1998; Cioce et al., 2006; Baldin et al., 2008), the soluble protein fraction was pre-extracted before fixation by treating the cells with 0.5%Triton X-100/PBS (Guillot et al., 2004). Analysis of images acquired with a widefield microscope suggested a potential co-localization between HP1β and PA28γ in some discrete areas of the nucleus (Fig. S4A, left panel, merged image and higher magnifications). Further analysis of HP1β and PA28γ proteins immunostaining, using a confocal microscope with Airyscan detection and image acquisition in Z-stacks followed by 3D reconstruction (Fig. S4B, left panel) suggested that indeed a small fraction of PA28γ co-localizes with HP1β with ~32 colocalization sites per nucleus in U2OS cells (Fig. S4B, right panel).

To strengthen this result, we used the *in situ* Proximity-Ligation Assay (*is*-PLA), which allows the detection of the close proximity between two proteins within cells (less than 40 nm, i.e. likely to be an interaction) (Soderberg et al., 2006). Due to the nuclear abundance of PA28γ and HP1β, we first verified the specificity of this approach by testing the signal between PA28γ and one of its known partners, the 20S proteasome, which is also highly abundant in the nucleus. Using antibodies raised against PA28γ and α4 (one subunit of the 20S proteasome), *is*-PLA revealed a characteristic dotted pattern throughout the nuclei of U2OS cells (Fig. S4C, upper panel). Quantification of the number of PLA dots per nucleus (see Materials and Methods) indicated less than 60 dots (Fig. S4C, bar graph), a number consistent with the low amount of α4/20S proteasome immunoprecipitated with PA28γ antibodies (see Fig. S2 and Jonik-Nowak et al., 2018), supporting the notion that this signal is specific. Then, using both PA28γ and HP1β antibodies (Fig. 4), *is*-PLA revealed on average 37 dots per nucleus (Fig. 4A, upper left panel and bar graph), a number in the same range as the number of co-localization sites (~32), as evidenced in Fig. S4B. Silencing of PA28γ expression with siRNAs (Fig. 4B), used as a negative control, abolished the PLA dots (Fig. 4A, lower panel and bar graph). Note that we also observed that a fraction of PA28γ colocalized in part with HP1α by the *is*-PLA approach (Fig. S4D). Taken together, these results indicate that a small fraction of PA28γ is in close physical proximity (and thus is likely to interact either directly or indirectly) to a fraction of the heterochromatin-binding protein HP1β.

**Figure 4.**
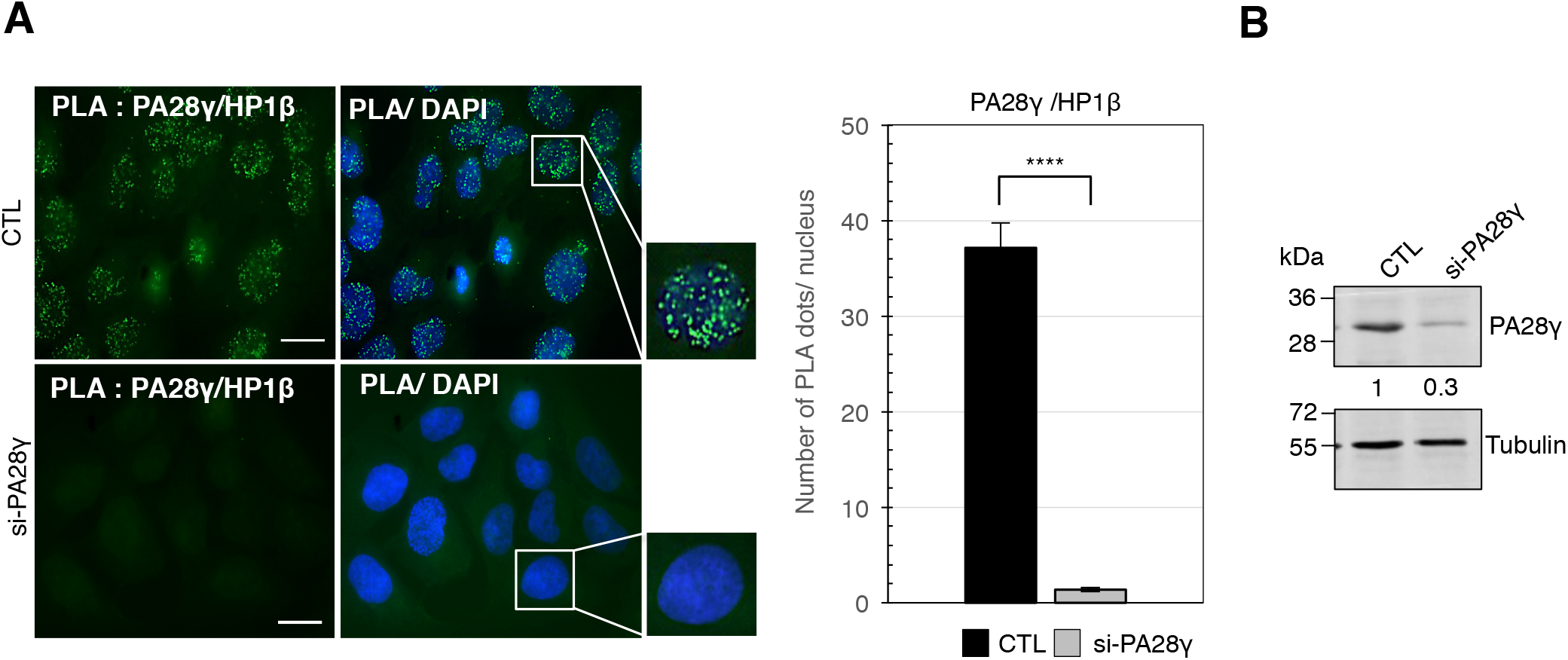
A fraction of PA28γ co-localizes with HP1β in U2OS cells. **A.** Control (CTL) or si-PA28γ treated U2OS cells were subjected to *is*-PLA using primary antibodies directed against HP1β and PA28γ, and DNA stained with DAPI. Positive PLA signals appear as green dots and higher magnification views of a nucleus are shown (left panel). Scale bars, 10 μm. Quantification of PLA dots was carried out using an ImageJ macro (see Materials and Methods). The number of PLA dots per nucleus for HP1β/PA28γ interaction in control (CTL) or si-PA28γ treated cells is shown graphically (right panel). Data represent the means ± SD from 3 independent experiments, the number of cells analyzed is *n* = 78 and *n* = 45 in control and si-PA28γ treated cells, respectively. The *p*-value was determined using Student’s *t*-test, **** (*p* = 0.0001). **B.** Immunoblot analysis of PA28γ expression level in total extracts of U2OS cells treated or not with si-PA28γ, used for *in situ* Proximity Ligation Assay (*is*-PLA). Tubulin was used as a loading control. The relative abundance of PA28γ (left panel) was quantified using ImageJ software.

### PA28γ is a chromatin compaction regulator as important as HP1 β

As PA28γ ensures chromatin compaction and partially co-localizes with HP1β in cells, we wondered whether PA28γ might play a role in chromatin compaction as important as HP1β. To address this question, we performed siRNA-mediated depletion of PA28γ, HP1β or both proteins in HeLa^H2B-2FPs^ cells (Fig. 5A) and compared the degree of chromatin compaction of these cells by FLIM-FRET approach (Fig. 5B,C). FRET measurements revealed a marked decompaction of chromatin upon PA28γ-knockdown that was even stronger than upon HP1β depletion (Fig. 5B). This decompaction was correlated with the clear disappearance of the most compacted states of the chromatin within nuclei (Fig. 5C, left panel). To complete these data, we extracted the FRET efficiency distribution curves related to the FRET efficiency map from individual nuclei (Fig. 5C, right panel). While siRNA-Luc only caused an increase in the high-FRET population (Fig. 5C right panel, blue curve) as compared to parental cells (Fig. 1D, right panel, blue curve), the quantitative analysis of the FRET distribution profiles revealed that PA28γ-knockdown by siRNA had a stronger effect than the PA28γ-knockout. This difference observed between PA28γ-knockdown and PA28γ-knockout might reflect potential compensatory mechanisms developed by the PA28γ-KO cell line to preserve cellular homeostasis and viability. Interestingly, as already observed in the Fig. 1, these results confirm that even in the presence of HP1 proteins the lack of PA28γ results in a strong decompaction of chromatin. This analysis also revealed a less pronounced decompaction of the chromatin, with some persistent high-FRET values remaining, upon si-HP1β depletion than after si-PA28γ depletion (Fig. 5C right panel, compare red and black curves). Most likely, the lower effect of HP1 β knockdown may be explained by the presence of the other HP1 isoforms. Furthermore, the depletion of both PA28γ and HP1β had no additional effect on chromatin decompaction compared to PA28γ alone (Fig. 5B and Fig. 5C right panel, compare green and black curves). Altogether these results strongly suggest that PA28γ is a key regulator of chromatin compaction as important as HP1β and that these two proteins might be involved in a similar regulatory pathway.

**Figure 5.**
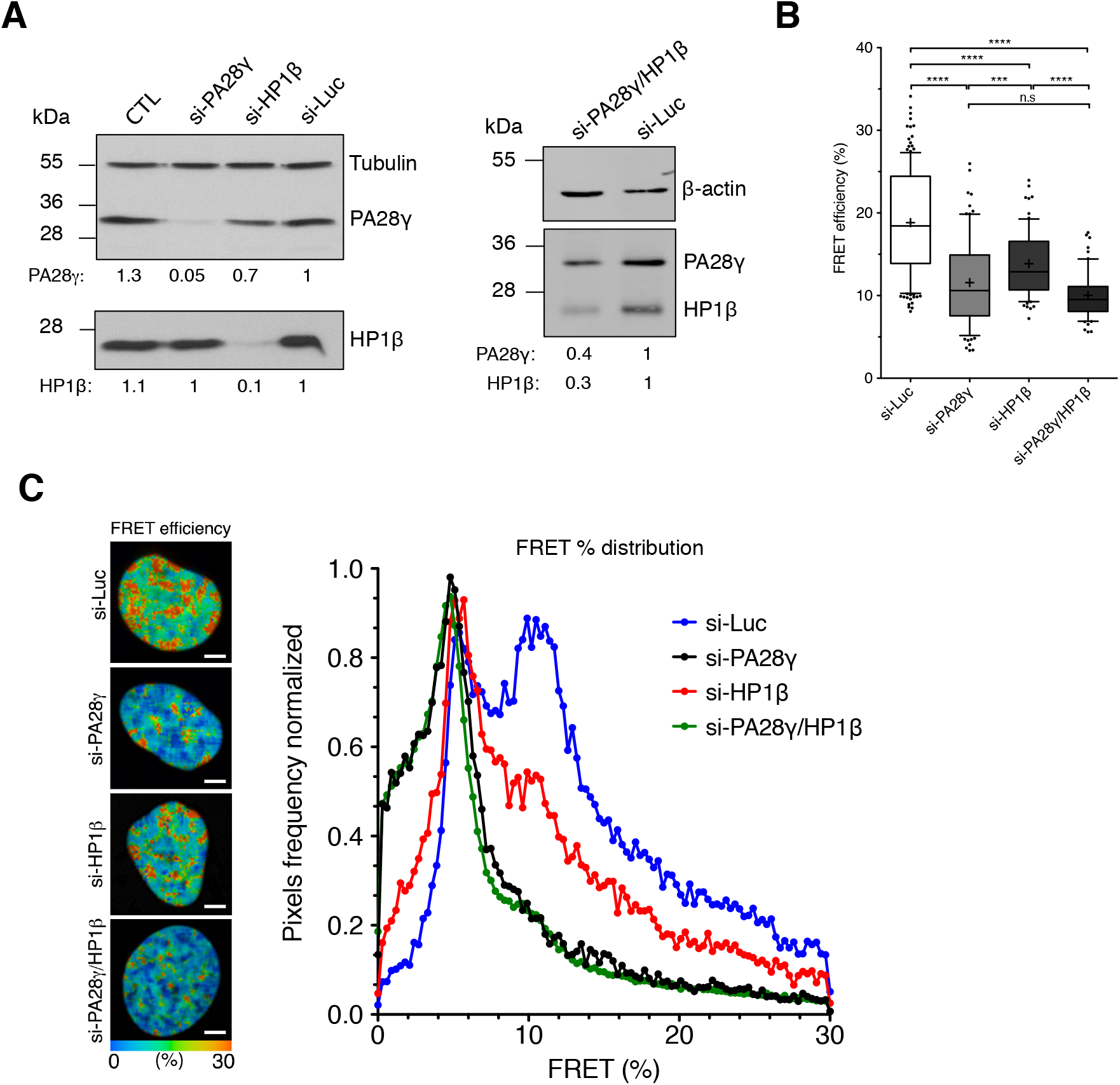
PA28γ is as crucial as HP1β for chromatin compaction. **A.** HeLa^H2B-2FPs^ cells (WT) were transfected with control si-Luc, si-PA28γ, si-HP1β or a mix of both siRNAs (si-PA28γ/HP1β) for 48 hours. Immunoblot analysis of PA28γ and HP1β protein levels in HeLa^2FPs^ following siRNA treatments were performed. Tubulin and anti-β actin antibodies were used as loading controls. The relative abundance of PA28γ and HP1β proteins was quantified using ImageJ software. **B.** Quantification of the mean FRET efficiencies were presented as box-and-whisker plots, where the thick line represents the median, the boxes correspond to the mean FRET values above and below the median, with the whiskers covering the 10-90 percentile range. Data represent the means ± SD from 4 independent experiments, the total number of analyzed cells is *n* = 152 (si-Luc), *n* = 85 (si-PA28γ), *n* = 73 (si-HP1β), *n* = 61 (si-PA28γ/HP1β). ns = not significant, *** *p* < 0.001, **** *p* < 0.0001 (Student’s *t*-test). **C.** Representative images of the spatial distribution of the FRET efficiency (percentage) in representative control si-Luc, si-PA28γ, si-HP1β, or both siRNAs (si-PA28γ/HP1β) treated HeLa^H2B-2FPs^ nuclei is depicted in a continuous pseudo-color scale ranging from 0 to 30% (left panel). Scale bars, 10 μm. Mean FRET distribution graph showing distinct populations of FRET efficiency in si-Luc (blue curve), si-PA28γ (black), si-HP1β (red), or both si-PA28γ/HP1β (green) treated HeLa^H2B-2FPs^ (right panel).

### PA28γ contributes to the maintenance of heterochromatin marks

Besides the key role of HP1 proteins, methylation of histone H3 on lysine 9 (H3K9me) (Maison and Almouzni, 2004; Grewal and Jia, 2007) and histone H4 on lysine 20 (H4K20me) (Schotta et al., 2004; Shoaib et al., 2018) have been shown to be important for maintaining the ground state of chromatin structure. We therefore set out to investigate whether PA28γ could regulate the chromatin compaction state through H3K9 and H4K20 methylation in cells. To achieve this, we first examined by western blot whether the loss of PA28γ might affect the steady-state levels of these epigenetic modifications. As shown in Fig. S5A, no significant change in the levels of H3K9me3 was observed in U2OS-KO-PA28γ cells compared to WT cells. By contrast, PA28γ depletion led to a decrease (~20%) in the steady-state level of H4K20me3 (Fig. S5B). This was accompanied by a significant decrease (~40%) in H4K20me1 (Fig. S5B), which is a pre-requisite for establishment of the H4K20me3 state (Tardat et al., 2007). These results led us to examine the variation of H3K9me3 and H4K20me3 at specific heterochromatin sequences. To this end, we carried out ChIP assays on parental (WT) and KO-PA28γ U2OS cells using antibodies against H3K9me3 and H4K20me3, and performed qPCR using the same primers as in Fig. 3E. We observed a significant decrease in H3K9me3 precipitation levels (≥50%) at the specific heterochromatin sequences (Fig. 6A) that was not detected by immunoblot analyses on total cell extract (Fig. S5A). Note that the difference observed between immunoblot and ChIP-qPCR assay could result from a difference of sensitivity of both techniques using H3K9me3 antibodies. Since H3K9me3 serves as a molecular anchor for HP1β, we checked its presence on these specific DNA sequences by ChIP-qPCR assay. Surprisingly, no obvious variation was observed for the sequences tested, except for the LINE-1 sequence (Fig. S5C), suggesting that either this decrease of H3K9me3 level is not sufficient to destabilize HP1β binding and/or the involvement of other HP1β domains, such as its chromoshadow domain (CSD) (Zeng et al., 2010; Liu et al., 2017; Kumar and Kono, 2020), would facilitate its binding when chromatin is decondensed. We also confirmed by ChIP-qPCR assay the substantial decrease of H4K20me3 (≥60%) on the same heterochromatin DNA sequences in KO-PA28γ *versus* WT U2OS cells (Fig. 6B).

**Figure 6.**
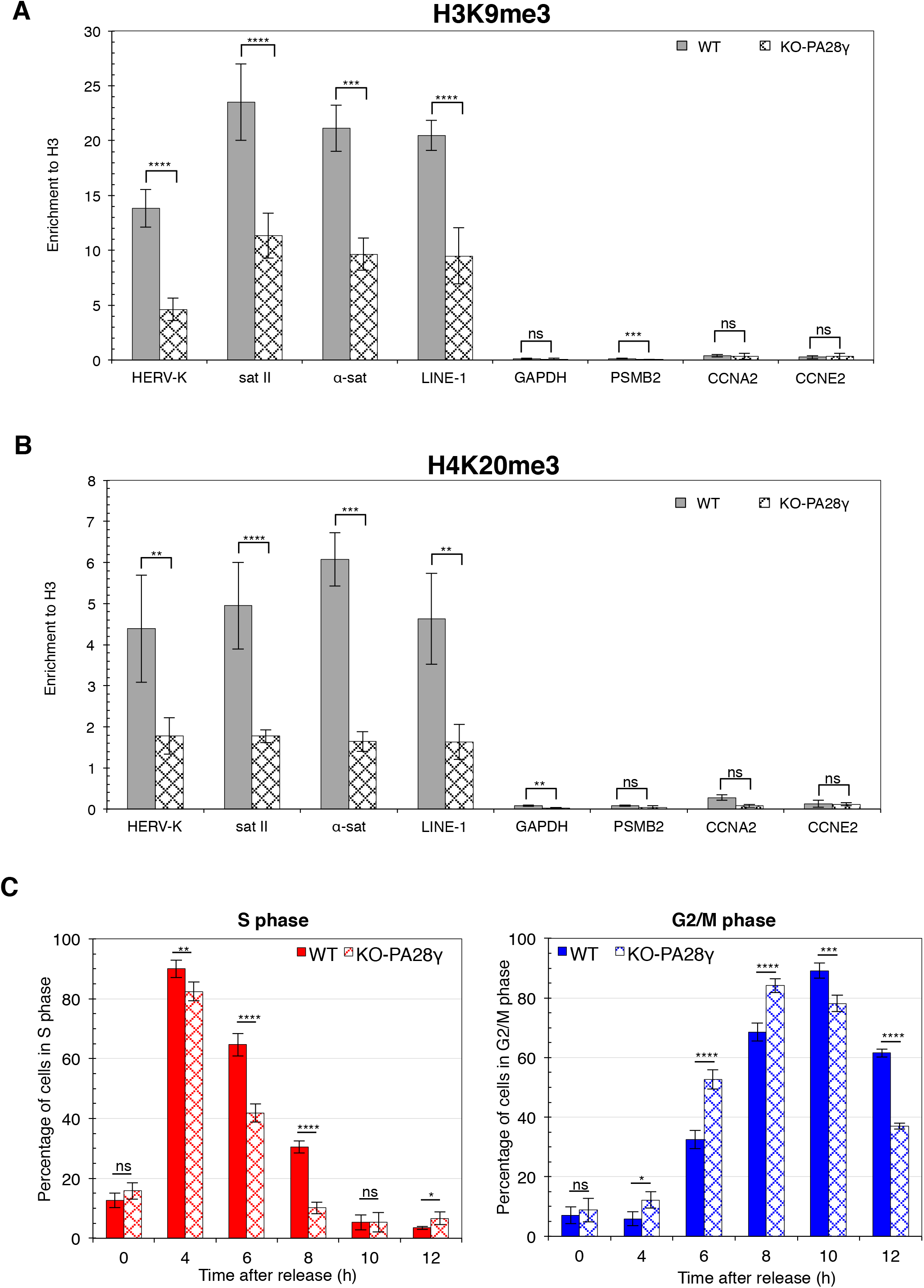
PA28γ contributes to the maintenance of heterochromatin marks and PA28γ depletion decrease S-phase duration. **A and B.** ChIP-qPCR analysis of H3K9me3 (A) and H4K20me3 (B) levels at different repetitive elements and gene promoters (as indicated on the x-axis) in WT *versus* KO-PA28γ U2OS cells. Data are represented as relative enrichments of each specific histone mark *versus* histone H3 control, as shown on the y-axis. Data are means ± SEM (*n* = 5 for H3K9me3 and histone H3, *n* = 3 for H4K20me3). Significance was calculated by Student’s *t*-test (ns = not significant, *\*p* < 0.05, *\*\*p* < 0.01, *\*\*\*p* < 0.001, *\*\*\*\*p* < 0.0001. *p*-values are presented in Table S1). **C.** Asynchronous parental (WT) and KO-PA28γ U2OS cells (AS), cells synchronized at the G1/S phase transition by a double thymidine block (0) and released for the times indicated were subjected to FACS analysis. Histograms representing the percentage of the cells in S (left panel) and G2/M (right panel) phases of the cell cycle are shown. Data represent the means ± SD from three biological repeats. *p* values were determined with a 2-way ANOVA for each time point. *p*-values are presented in Table S3.

We also investigated whether the loss of PA28γ induced a change of H3K4me3, a modification considered as an epigenetic biomarker of transcription activation (Howe et al., 2017). No significant variation in H3K4me3 levels was detected by immunoblot analyses on total cell extracts (Fig. S5D). This absence of variation was confirmed by ChIP-qPCR assay using primers on which a significant decrease of H3K9me3 and/or H4K20me3 was observed (Fig. S5E), suggesting that PA28γ depletion has no significant impact on the transcription of the sequences tested. These results are in line with previous data showing that PA28γ knockdown has no impact on the global transcription level (Cioce et al., 2006; Baldin et al., 2008), and our results indicating no variation in the transcription of the heterochromatin DNA sequences by RT-qPCR (data not shown).

The importance of H4K20me1/me3 and chromatin compaction in cell cycle progression and in the regulation of DNA replication (Brustel et al., 2017; Shoaib et al., 2018) prompted us to examine whether the loss of PA28γ might impact on cell cycle progression. To this end, parental (WT) and PA28γ-depleted (KO-PA28γ) U2OS cells were synchronized with a double thymidine block and then released from the G1/S transition before analysis for cell cycle progression by DNA content analysis using flow cytometry (Fig. 6C). Our data indicated that cells lacking PA28γ led to early S-phase as parental U2OS cells, but progressed faster and exit S-phase earlier compared to the wild-type U2OS cells (Fig. 6C, left panel). Consistent with this, KO-PA28γ cells showed an earlier entrance into G2 phase (Fig. 6C, right panel). This shortening of S phase (~1 hour) in KO-PA28γ cells was confirmed by immunoblotting using cell cycle markers including cyclin E (a marker of G1 to S-phase transition) and the phosphorylation of histone H3 on serine 10 (a mitosis marker) (Fig. S6). Altogether, these results suggest that the chromatin decompaction and alterations in the levels of heterochromatin histone marks upon loss of PA28γ are not toxic *per se* but accelerate S-phase progression, likely by favouring accessibility of the most-compact chromatin regions to the replication machinery.

## Discussion

This study provides several pieces of evidence that PA28γ, which is known as a nuclear activator of the 20S proteasome, is also an essential regulator of chromatin structure.

We demonstrate that PA28γ plays a key role in the process of chromatin compaction by showing that the depletion of PA28γ, by knockout and/or knockdown approaches: i) induces a decompaction of the highly structured fraction of the chromatin, even in the presence of HP1 proteins, as visualized in living cells with our quantitative chromatin compaction assay, and ii) causes the decompaction of *lacO* DNA repeats integrated into a pericentromeric heterochromatin domain. As summarized in Fig. 7, we show that PA28γ is present on chromatin regions enriched for HP1β and contributes to the maintenance of heterochromatin features, such as H3K9 and H4K20 tri-methylation.

**Figure 7.**
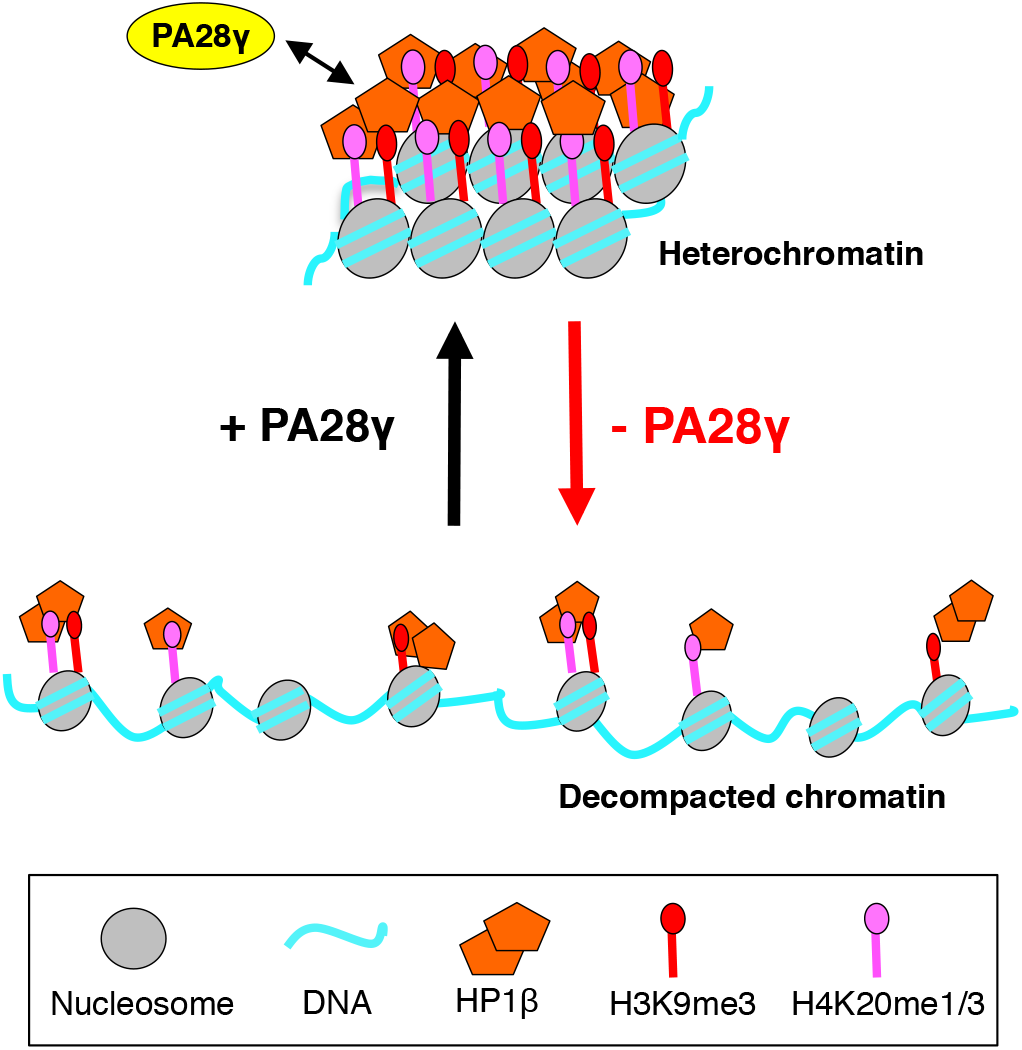
Schematic summary of findings. Our results show that PA28γ, as HP1β, is associated with heterochromatin domains. In the absence of PA28γ, the tri-methylation level of H3K9 and H4K20 is substantially reduced, as a consequence preventing the compaction of these heterochromatin domains, even in the presence of HP1β.

A striking result of our study is that the chromatin structural role of PA28γ and its impact on the compaction of heterochromatin is as important as HP1β, which is considered as a key regulator of heterochromatin domains and maintenance. We find that a small fraction of PA28γ co-localizes with HP1β in the nucleus, and the difficulty to detect this co-localization suggests that it could occur in dynamic and/or transient structures. Interestingly, recent studies show that HP1 proteins have the capacity to form liquid-like droplets (also called condensates) resulting from a liquid-liquid phase separation (LLPS) mechanism (Larson et al., 2017; Strom et al., 2017). This property facilitates the enrichment of transient complexes that could be rapidly assembled and disassembled, and the exchange of various proteins required for heterochromatin compaction. Although recent results suggest that heterochromatin maintenance is independent of liquid droplet formation of HP1α in mouse chromocenters and rather involves collapsed chromatin globules, HP1 proteins form transient droplets in the cells that could participate in the structure of chromatin subcompartments (Erdel et al., 2020). Considering this point, it is important to underline the fact that PA28γ is detected in various membraneless compartments such as NS, CB and PML bodies, considered to be liquid-like protein droplet organelles (Erdel and Rippe, 2018; Sawyer et al., 2019). Although the mechanism by which PA28γ is recruited into these nuclear bodies still remains to be unraveled, the interaction of PA28γ with specific proteins present in these condensates plays a crucial role in the control of their dynamics (Cioce et al., 2006; Zannini et al., 2009; Jonik-Nowak et al., 2018). For example, the interaction of PA28γ with Chk2, a cell cycle checkpoint kinase that localizes in PML bodies, is required for the control of PML body number (Zannini et al., 2009). In this context, the capacity of HP1 proteins to form condensates could participate in the transient enrichment of PA28γ in specific domains of the chromatin that might facilitate the establishment of PA28γ interaction with proteins required for chromatin compaction.

How PA28γ, which has no known enzymatic activity, could favor proper maintenance of chromatin structure is still an open question. Our results suggest that the function of PA28γ function in chromatin compaction is likely independent of its proteasome-regulatory function, since a PA28γ mutant deleted from its C-terminal portion still promotes chromatin compaction. Indeed, the binding of the C-terminal extremity of PA28 activators to the 20S α-ring is the first essential step for complex formation and activation of the proteasome (Förster et al., 2005). Although we cannot at this stage exclude the possibility of a transient interaction of PA28γ mutant with the 20S proteasome in cells, a direct regulation of chromatin compaction by a PA28γ-dependent proteolysis event seems unlikely. Therefore, since PA28γ depletion induces a significant decrease of H3K9me3 (≥50%), H4K20me1 (≥40%) and H4K20me3 (≥60%), it is conceivable that PA28γ acts either by facilitating the function of the lysine methyltransferases Suv39h, PR-Set7 or Suv4-20h responsible for H3K9 trimethylation, H4K20 monomethylation and H4K20 trimethylation, respectively, or by inhibiting specific histone demethyltransferases, or other proteins complexes involved in chromatin remodeling. It is interesting to note that the PA28γ interactome contains two major interactors, BRD9 and SMARCA4 (BRG1) (Jonik-Nowak et al., 2018), which are two subunits of a newly-defined ATP-dependent chromatin remodeling complex (Alpsoy and Dykhuizen, 2018). However the physiological significance of these interactions in the new function of PA28γ in chromatin compaction remains to be determined.

Chromatin alterations occurring upon loss of PA28γ neither impact cell viability nor induce a strong phenotype, as observed upon HP1β depletion in MEF cells (Bosch-Presegue et al., 2017) or in HP1-triple knockout in hepatocytes (Saksouk et al., 2020). However, our data reveal a change in cell cycle progression with a decrease in S-phase duration, suggesting that the accessibility and/or the progression of the replication machinery could be facilitated by the decompaction of the most condensed chromatin domains.

It is noteworthy that previous studies reported the consequences of PA28γ depletion on chromatin-related processes, such as centrosome maintenance and chromosomal stability (Zannini et al., 2008) and DNA repair (Levy-Barda et al., 2011). Our present observations suggest that the role of PA28γ in the regulation of chromatin structure could be the common mechanism that links these processes to PA28γ. Indeed, alterations in H3K9 methylation, as observed in KO-PA28γ cells, results in an increase in chromosome segregation errors, which have been linked to a role of pericentromeric heterochromatin in the proper assembly of centromeres (Peters et al., 2001; Peng and Karpen, 2009). The reported increase of aneuploidy under PA28γ-knockdown (Zannini et al., 2008) could also result from the decrease of H3K9me3 observed in our study. In the same vein, PA28γ depletion did not spontaneously induce DNA damage, but leads to an increase of cellular radiomimetic sensitivity and a substantial delay in DNA double-strand-break (DSBs) repair (Levy-Barda et al., 2011). This effect could also result from the contribution of PA28γ towards maintaining appropriate levels of H3K9me3 and H4K20me1/3 since these histone modifications have been involved in promoting or inhibiting the recruitment of specific repair proteins, which directly affect DNA damage repair efficiency (Price and D’Andrea, 2013).

Altogether, our data reveal that PA28γ is a novel and crucial factor in the regulation of chromatin compaction. Although much remains to be understood regarding its exact contribution to this process, our findings undoubtedly open new avenues of research for a deeper understanding of the complex mechanisms that control chromatin organization.

## Materials and methods

### Plasmids

For Cas9-mediated gene disruption, guide RNA (GGAAGTGAAGCTCAAGGTAGCGG) targeting PA28γ *(PSME3)* was selected using ChopChop (https://chopchop.cbu.uib.no/) and oligonucleotides were subcloned into pMLM3636 (a gift from Keith Joung, Addgene plasmid #43860) and pUC57-U6 (a gift from Dr. E Bertrand’s laboratory, IGMM, Montpellier, France). For rescue experiments, PA28γ ORF WT or minus the C-terminal 14 amino acids (ΔC) were cloned into pSBbi-Pur (a gift from E. Addgene plasmid #60523) according to (Kowarz et al., 2015). The resulting vector was co-transfected with pCMV(CAT)T7-SB100 (gift from Zsuzsanna Izsvak, Addgene plasmid #34879) into recipient cells, and puromycin-resistant single colonies were selected for re-expression of PA28γ WT or ΔC proteins. pEGF-LacI (Jegou et al., 2009) was a generous gift from Prof. K. Rippe (DKFZ, Heidelberg, Germany).

### Antibodies

The following antibodies were used at 1:1000 dilution for immunoblotting and 1-3 μg/ml for immunoprecipitation: anti-PA28γ (rabbit polyclonal BML-PW8190, ENZO Life Sciences), anti-α4 (1:2000; mouse monoclonal BML-PW8120, ENZO Life Sciences); anti-PA28γ (mouse monoclonal, 611180, BD Transduction); anti-HP1α (rabbit polyclonal, 2616S, Cell Signaling); anti-HP1β (rabbit monoclonal (D2F2), 8676S, Cell Signaling and mouse monoclonal (1MOD-1A9) 39979, Active Motif); anti-GFP (mouse monoclonal, Clone 7.1, 11814460001, ROCHE, Sigma); anti-RFP (rat monoclonal, 5F8, Chromotek); anti-β-actin (rabbit monoclonal, 13E5, Cell Signaling); anti-H3K9me3 (mouse monoclonal, clone 2AG-6F12-H4, 39285, Active Motif); anti-H3K4me3 (mouse monoclonal (clone 2AG-6F12-H4) 39285, Active Motif); anti-histone H3 (rabbit polyclonal, ab1791, Abcam); anti-H4K20me1 (rabbit polyclonal, #9724, Cell Signaling Technology); anti-H4K20me3 (rabbit monoclonal, #5737, Cell Signaling Technology); anti-histone H1 (rabbit polyclonal, PA5-30055, Thermo Fisher) anti-α-tubulin (mouse monoclonal, T9026, Sigma-Aldrich, 1:6,000). Fluorescent secondary antibodies conjugated either to Alexa Fluor 488 or 594 (1:1,000), or to DyLight 680 or 800 (1:10,000) were purchased from Thermo Fisher Scientific. Secondary antibodies conjugated to HRP were purchased from Bio-Rad SA (1:10,000).

### Cell culture, transfections, cell synchronization and FACS analysis

U2OS (HTB-96) cells, obtained from ATCC, were grown in DMEM (Lonza) containing 4.5 g/L glucose, 10% heat-inactivated fetal bovine serum (Biowest), 2 mM glutamine, 100 U/ml penicillin and 10 μg/ml streptomycin (Lonza). U2OS-LacO (F42B8) cells (a generous gift of Prof. K. Rippe, DKFZ, Heidelberg, Germany) were grown in the same media as U2OS but containing G418 (500 μg/ml) (Jegou et al., 2009). Establishment and characterization of parental HeLa^H2B-GFP^ and HeLa^H2B-2FPs^ (H2B-GFP and mCherry-H2B) cell lines were previously described (Lleres et al., 2009). Of note: after thawing, cells were cultured for one week before seeding, for all experiments.

For transient PA28γ and HP1β knockdown experiments, U2OS-LacO and/or HeLa (H2B-GFP or 2FPs) cells were transfected with 20 nM Luciferase targeting siRNA (si-Luc, 5’-CGTACGCGGAATACTTCGA-3’) used as negative control, or -PA28γ *(PSME3),* and – HP1β *(CBX1)* targeting siRNA (si-PA28γ: 5’-GAAUCAAUAUGUCACUCUA-3’; si-HP1β: 5’-AGGAAUAUGUGGUGGAAAA-3’) purchased from Eurofins Genomics, using Lipofectamine RNAiMAX (Thermo Fisher Scientific) and examined after 2 days. Where indicated, cells were transiently transfected with 0.5 μg/ml DNA using JetPEI™ (Ozyme), according to the manufacturer’s instructions and analyzed after one day. Stable U2OS (Jonik-Nowak et al., 2018), HeLa^H2B-GFP^- and HeLa^2FPs^-KO-PA28γ cell lines were generated by cotransfection of PSME3/PA28γ sgGuide and pX459 vectors (a gift from Feng Zhang, Addgene plasmid #62988), and cells were selected with puromycin (1 μg/ml). Single clones were then expanded and analyzed by western blotting using PA28γ antibodies. Synchronization of cells at the G1/S phase transition was performed by double thymidine block as described (Thomas et al., 2014). For Fluorescence-Activated Cell Sorting (FACS) analysis, cells were fixed with 70% ethanol and conserved at −20°C. Before analysis, cells were washed with PBS, resuspended in PBS containing RNAse A (1 mg/ml, Sigma-Aldrich) and propidium iodide (10 μg/ml, Sigma-Aldrich) and incubated for 30 min at room. Samples were run on a FACSCalibur Cell Analyzer (Becton-Dickinson), and data analysis was performed using CellQuest Pro software (Beckton-Dickinson).

### Immunofluorescence and is-PLA assays

Cells on coverslips were fixed in 3.7% paraformaldehyde/PBS at room temperature, then permeabilized with 0.25% Triton X-100 in PBS for 5 min, followed by incubation in methanol (100%) at −20°C for 10 min. After washes with PBS, cells were blocked with 1% Calf Serum/PBS for 15 min. Incubation with primary antibodies (anti-PA28γ 1:6,000 for BML-PW8190 or 1:1,000 for 611180); anti-α4 (1:4,000 BML-PW8120); anti-HP1α (1:800, 2616S); anti-HP1β (1:1,000 8676S and 1MOD-1A9)) was carried out at 37°C for 1 hour in a humidified atmosphere. After washes, cells were incubated with Alexa Fluor conjugated secondary antibodies for 40 min at RT. DNA was stained with 0.1 μg/ml DAPI (4’,6-diamidino-2-phenylindole, dihydrochloride, Sigma-Aldrich) solution 5 min at RT, cells were washed twice in PBS and finally once in H_2_O. Coverslips were mounted on glass slides using ProLong Gold anti-fade reagent (Thermo Fisher Scientific). For *in situ* proximity ligation assays (*is*-PLA), cells on coverslips were fixed and permeabilized as above. Coverslips were then blocked in a solution provided by the Duolink^®^ kit (Sigma-Aldrich). Cells were then incubated with antibodies as described above. Duolink^®^ *In Situ* PLA Probe Anti-Rabbit MINUS and Anti-Mouse PLUS and Duolink^®^ *In Situ* Detection Reagents (Sigma-Aldrich) were used, according to the manufacturer’s instructions. In some specific experiments, cells were permeabilized prior to fixation with 0.5% Triton X-100 in PBS for 5 min at 4°C for the co-localization between endogenous PA28γ and HP1β in U2OS cells.

2D and Z-stack images were acquired with 63x/1.32 NA or 100x/1.4 NA oil immersion objective lenses using a DM 6000 microscope (Leica). Microphotographs were taken with a 12-bit CoolSNAP HQ2 camera. Images were acquired as TIFF files using MetaMorph imaging software (Molecular Devices). For quantitative analyses of PLA dots, Z-stacks were acquired every 0.3 μm (Z step) with a range of 6-7.5 μm. For endogenous detection, images (as a Z stack, slices every 200 nm) were also acquired on a Zeiss LSM 880 point scanning confocal microscope equipped with a 63x Plan-Apochromat 1.4NA oil immersion objective (Zeiss) and using the 488 nm and 561 nm laser lines with the Airyscan detector. The Zeiss Zen black software was used to process the Airyscan raw Images. Co-localization in 3D, between PA28γ and HP1β, was analyzed using the Imaris (Bitplane) co-localization module.

The number of PLA-dots and the size of GFP-LacI dots were determined using ImageJ (1.49v). Custom macros were created to automatically quantify these different parameters. The script allows the creation of a mask of DAPI image to isolate the nucleus of each cell and create a maximum intensity projection (MIP) of the Z-stacks or the image. The mask is used in the MIP to count the number of PLA-dots of each nucleus via an appropriate threshold. The “Analyze Particles” tool of ImageJ was used to calculate the size of each GFP-LacI dot.

### FLIM-FRET Microscopy

FLIM-FRET data were acquired with a Zeiss LSM 780 laser scanning microscope coupled to a 2-photon Ti:Saphire laser (Chameleon Ultra II tunable 680–1080 nm, Coherent) producing 150-femtosecond pulses at 80 MHz repetition rate and a Time Correlated Single Photon Counting (TCSPC) electronics (SPC-830; Becker & Hickl GmbH) for time-resolved detection. Enhanced green fluorescent protein (EGFP) and mCherry fluorophores were used as a FRET pair. The two-photon excitation laser was tuned to 890 nm for selective excitation of the donor fluorophore. The LSM780 microscope is equipped with a temperature-and CO_2_- controlled environmental black wall chamber. Measurements were acquired in live cells at 37°C, 5% CO_2_ with a 63x/1.4 oil Plan-Apochromat objective lens. A short-pass 760-nm dichroic mirror was used to separate the fluorescence signal from the laser light. Enhanced detection of the emitted photons was afforded by the use of the HPM-100 module (Hamamatsu R10467-40 GaAsP hybrid PMT tube). The FLIM data were processed using SPCimage software (Becker & Hickl GmbH).

### FLIM-FRET analysis

FLIM-FRET experiments were performed in HeLa cells stably expressing H2B-GFP alone (HeLa^H2B-GFP^) or with mCherry-tagged histone H2B (HeLa^H2B-2FPs^). 5 x 10^4^ cells were seeded in a FluoroDish 35 (FD35-100, World Precision Instruments). For siRNA experiments, 24 hours after seeding, cells were transfected with 20 nM of siRNA (against Luciferase, PA28γ or HP1β) and FLIM-FRET experiments were performed 48 hours later. 30 min prior to imaging, the culture medium was changed to complete DMEM medium without phenol red. An acquisition time of 90 s was set up for each FLIM experiment. The analysis of the FLIM measurements was performed by using SPCImage software (Becker & Hickl, GmbH). Because FRET interactions cause a decrease in the fluorescence lifetime of the donor molecules (EGFP), the FRET efficiency was calculated by comparing the FLIM values obtained for the EGFP donor fluorophores in the presence (HeLa^H2B-2FPs^) and absence (HeLa^H2B-GFP^) of the mCherry acceptor fluorophores. FRET efficiency (E FRET) was derived by applying the following equation:

E FRET = 1-(τDA / τD) at each pixel in a selected region of interest (nucleus) using SPCImage software. τDA is the mean fluorescence lifetime of the donor (H2B-EGFP) in the presence of the acceptor mCherry-H2B in HeLa^H2B-2FPs^ cells and τD is the mean fluorescence lifetime of H2B-EGFP (in the absence of acceptor) in HeLa^H2B-GFP^ cells. The FRET distribution curves from nuclei were displayed from the extracted associated matrix using SPCImage and then normalized and graphically represented using Microsoft Excel and GraphPad Prism software. For each experiment, FLIM was performed on multiple cells from several independent experiments (see Figure legends).

### Immunoprecipitation and immunoblotting

For immunoprecipitation, cells were lysed in lysis buffer (50 mM Tris-HCl, pH 7.5, 150 mM NaCl, 5 mM MgCl_2_, 1% IGEPAL CA-630, 0.5% sodium deoxycholate (DOC), 0.1% SDS, 1 mM DTT, 5 mM EDTA, 50 mM NaF and 1 mM Na_3_VO_4_) in the presence of complete EDTA-free protease inhibitor cocktail (Roche Life Science) for 20 min at 4°C. Lysates were clarified by centrifugation for 10 min at 10,000 x g and the protein concentration of the supernatant was determined using BSA as a standard (CooAssay protein dosage reagent, Interchim). Total lysate (200 μg) was pre-cleared for 30 min, and immunoprecipitations were performed using the antibodies indicated and protein A magnetic beads (Dynal, Lake Success, NY) for 2 hours at 4°C with constant gentle stirring. After several washes, bead pellets were boiled in 2x Laemmli buffer, separated by SDS-PAGE, and subjected to immunoblotting.

### ChIP-qPCR

ChIP experiments with U2OS were performed as described previously (Brustel et al., 2017). Briefly, cells were fixed with 1% formaldehyde (10 min) and quenching was performed with 125 mM Glycine. After a PBS wash, cells were resuspended in buffer A (10 mM Tris-HCl pH 8, 10 mM KCl, 0.25% Triton X-100, 1 mM EDTA, 0.5 mM EGTA) for 5 min on ice. After centrifugation, nuclei were extracted with buffer B (10 mM Tris pH 8, 200 mM NaCl, 1 mM EDTA, 0.5 mM EGTA) for 10 minutes on ice. To extract chromatin, nuclei were resuspended in Lysis buffer (10 mM Tris-HCl pH 8, 140 mM NaCl, 0.1% SDS, 0.5% Triton X-100, 0.05% DOC, 1 mM EDTA, 0.5 mM EGTA). After sonication with an EpiShear probe sonicator (Active Motif) to obtain chromatin fragments less than 800 bp, ChIP was performed with 15-30 μg of sheared chromatin incubated with protein A magnetic beads (Invitrogen, Thermo Fisher) coupled with the appropriate antibody, as follows: anti-H3pan (1 μl/ChIP, C15310135 Diagenode), anti-H3K9me3 (2 μl/ChIP, C15410056 Diagenode), anti-H3K4me3 (1 μl/ChIP, C15410003 Diagenode) anti-H4K20me3 (2 μl/ChIP, C15410207, Diagenode), anti-PA28γ (0.5 μl /ChIP, ENZO Life Sciences), anti-HP1β (2 μl/ChIP, rabbit monoclonal (D2F2), 8676S, Cell Signaling). ChIP experiments were performed at least three times from independent chromatin preparations and quantitative PCR analyses of ChIP DNAs were performed using a SYBR green quantitative PCR kit (Invitrogen, Thermo Fisher) and a LightCycler 480 II instrument (Roche) under conditions standardized for each primer set. The amount of DNA in ChIP samples was extrapolated from standard curve analysis of chromatin DNA before immunoprecipitation (input), and values were represented as the ratio between the percentage of input obtained for each antibody to the ones obtained for histone H3. Primer sets used for qPCR: HERV-K For 5’-TGCCAAACCTGAGGAAGAAGGGAT-3’ and HERV-K Rev 5’-TGCAGGC ATTAAACATCCTGGTGC-3’, Sat-II For 5’-CCAGAAGGTAATAA GTGGCACAG-3’ and Sat-II Rev 5’-CCCTCCTTGAGCATTCTAACTACC-3’, α-Sat For 5’-GAAACACTCTTTCTGCACTAC CTG-3’ and α-Sat Rev 5’-GGATGGTTCAACACT CTTACATGA-3’ (Djeghloul et al., 2016), LINE-1 5’UTR For 5’-CAGCTTTGAAGA GAGCAGTGG-3’ and LINE-1 5’UTR Rev 5’-GTCAGGGACCCACTTGAGG-3’ (Filipponi et al., 2013), CCNA2 For 5’-ACTAGACGTCCCAGAGCTAAA-3’ and Rev, 5’-TGTCCGAAGGCTGACTCTAA-3’, CCNE2 For 5’-AAGCGTTAGAAATGGCAGAAAG-3’ and CCNE2 Rev 5’-TCTCTCCCTAATTTACCTGTAGGA-3’, GAPDH For 5’-GCACGTAGCTCAGGCCTCA AGAC-3’ and GAPDH Rev 5’-GACTGTCGAA CAGGAGGAGCAGAG-3’ and PSMB2 For 5’-GTGCTTGTCTCTGGGATCGT-3’ and PSMB2 Rev 5’-AAACTGGGCGTCACATA AGG-3’ (https://www.chipprimers.com/).

### Statistics

Error bars represent standard deviations unless otherwise noted. Different tests were used to determine significance, and as noted in the legend. Differences were considered significant when P < 0.05 and are indicated by different numbers of asterisks, as follows: *, *p* < 0.05; **, *p* < 0.01; ***, *p* < 0.001; and ****, *p* < 0.001.

## Author contributions

VB conceived the project and supervised the study. DF, DL, CG, SB, CBA, and VB performed experiments and analyzed data. FM generated reagents. OC provided conceptual advice on the study and the interpretation of the data. CBA and VB wrote the article with input from all of the authors.

## Acknowledgements

We thank K. Rippe (DKFZ, Heidelberg, Germany) for providing U2OS-LacO (F42B8) cell and the pEGFP-LacI vector, P. Fort for help with statistical analysis, N. Morin for help with Airyscan microscopy and E. Julien for useful scientific discussions and advice; and the Montpellier Ressources Imagerie (MRI) platform, a member of the National Infrastructure France-BioImaging supported by the French National Agency (ANR-10-INSB-04, Investments for the Future). We thank J. Hutchins for checking the scientific English.

## Supplementary Figures

**Figure S1. PA28γ -depletion does not alter the expression level of H2B-GFP or mCherry-H2B.**

**A.** Immunoblot analysis of H2B-GFP and mCherry-H2B expression level in total extracts from parental (WT) and KO-PA28γ HeLa^H2B-2FPs^ cells (left panel). Tubulin was used as a loading control. The relative abundance of H2B proteins was quantified using ImageJ software. Graphical representation of the relative abundance of H2B-GFP and mCherry-H2B, detected with an anti-GFP and anti-RFP, respectively, and normalized to tubulin (right panel). The mean ± SD is from four independent experiments. Statistical significance was evaluated based on Student’s *t*-test, ns = not significant *(p* = 0.2027 and 0.4024 for H2B-GFP and mCherry-H2B, respectively).

**B.** Quantification of the H2B-GFP and mCherry-H2B fluorescence intensities in WT and KO-PA28γ HeLa^H2B-2FPs^ cells. The total number of cells analyzed is *n* = 172 (WT), *n* = 183 (KO-PA28γ). Statistical significance was evaluated with Student’s *t*-test, ns= not significant.

**C.** FRET analysis in WT, KO-PA28γ HeLa^H2B-FPs^ cells, and WT HeLa^H2B-FPs^ cells treated with Trichostatin A (TSA, 200ng/ml, 24 h). The statistical analysis of the mean FRET efficiency percentage is presented as box-and-whisker plots. The thick line represents median, the boxes correspond to the mean FRET values above and below the median, with the whiskers covering the 10-90 percentile range. The total number of nuclei analyzed is *n* = 154 (WT), *n* = 132 (KO-PA28γ), and *n* = 33 (WT + TSA), **** *p* < 0.0001 (Student’s *t*-test).

**Figure S2. PA28γ -Δ C-mutant does not interact with the 20S proteasome.**

Whole-cell extracts from parental HeLa^H2B-2FPs^ (WT), PA28γ-knockout (KO-PA28γ) cells and KO cells re-expressing the wild-type (KO/KI-WT#8) form or the ΔC-mutant (KO/KI-ΔC) of PA28γ were subjected to immunoprecipitation using anti-PA28γ antibodies. Immunoblots of the pull-down (IP-PA28γ) and the supernatant (SN-IP, 1/10^eme^) from whole-cell extracts were probed with the antibodies indicated.

**Figure S3. PA28γ -depletion does not affect H1, H3 or HP1β expression level and HP1β is present at the same repetitive elements than PA28γ.**

**A.** Immunoblot of whole-cell extract (30 μg) from asynchronous parental (WT) and KO-PA28γ (KO-PA28γ) U2OS cells, using anti-PA28γ. Tubulin was used as a loading control.

**B.** Immunoblot analysis of histone H3 and H1 expression level in total extracts from WT and KO-PA28γ U2OS cells (left panel). Tubulin was used as a loading control. The relative abundance of histone H3 and H1 proteins was quantified using ImageJ software. Graphical representation of the relative abundance of histone H3 and H1 normalized to tubulin and histone H3, respectively (right panel). The mean ± SD is from four independent experiments. Statistical significance was evaluated based on Student’s *t*-test, ns = not significant. (*p* = 0,7560 and 0. 0,92033 for H3 and H1, respectively).

**C.** Immunoblot analysis of HP1β expression level in total extracts from WT and KO-PA28γ U2OS cells (left panel). Histone H3 was used as a loading control. Graphical representation of the relative abundance of HP1β normalized to histone H3 (right panel). The mean ± SD is from three independent experiments. Statistical significance was evaluated based on Student’s *t*-test, ns = not significant (*p* = 0,99619).

**D.** Immunoblot analysis of HP1β expression level in total extracts from U2OS cells treated or not with si-HP1β (left panel). Tubulin was used as a loading control. The relative abundance of HP1β proteins was quantified using ImageJ software. ChIP-qPCR analysis of HP1β levels at different repetitive elements (as indicated on the x-axis) in U2OS cells treated with si-Luc or si-HP1β. Data are represented as relative enrichment of HP1β *versus* histone H3 control, as shown on the y-axis (right panel). Data are means +/− SEM (*n* = 5). Significance was calculated using Student’s *t*-test, ns = not significant, * *p* < 0.05, ** *p* < 0.01 and *** *p* < 0.001. *p*-values are presented in Table S2.

**Figure S4. Co-localization of a fraction of PA28γ with HP1β and HP1α.**

**A.** Asynchronously-growing wild-type (left panel) and KO-PA28γ (right panel) U2OS cells were pre-permeabilized with 0.5% Triton-X100 to extract soluble proteins before fixation and the detection of endogenous HP1 β and PA28γ by indirect immunofluorescence using anti-HP1β and PA28γ antibodies. Representative merged images of HP1β (green) and PA28γ (red) are shown (right panels), higher-magnification views are shown for U2OS-WT cells. Scale bars, 10 μm.

**B.** A representative Airyscan confocal Z-projected image showing the co-detection of HP1β (green) and PA28γ (red) (left) in U2OS cells treated as in A. Co-localizations of both proteins along the cross are shown (left panel). Scale bars, 5 μm. Using the co-localization module of Imaris, a representative image of HP1β (green), PA28γ (red) corresponding to a 3D image (middle panel) is shown with the corresponding image showing only co-localization spots (white/grey, right panel). Scale bars, 5 μm.

**C.** *In situ* proximity ligation assay (*is*-PLA) was carried out in asynchronous U2OS cells using primary antibodies directed against PA28γ (rabbit polyclonal) and the α4 subunit of the 20S proteasome (mouse monoclonal) (CTL) or with α4 and without PA28γ antibodies (w/o anti-PA28γ) and DNA was stained with DAPI. Positive PLA signals appear as green dots and higher magnification views of a nucleus are shown (left panel). Scale bars, 10 μm. The number of PLA dots per nucleus in cells treated with both antibodies (CTL) or with only α4 antibodies (w/o anti-PA28γ) is shown on the bar graph (right panel). Data represent the mean ± SD from 3 independent experiments, the number of cells analyzed was *n* = 38 and *n* = 42 in control cells and cells treated without primary PA28γ antibody, respectively. The *p*-value was determined using Student’s *t*-test, \*\*\*\**p* ≤ 0.0001.

**D.** *Is*-PLA was carried out in U2OS cells using primary antibodies directed against PA28γ (mouse monoclonal) and HP1α (rabbit polyclonal) (CTL) or with PA28γ and without HP1α antibodies (w/o anti-HP1α) and DNA was stained with DAPI. Positive PLA signals appear as green dots and a higher magnification view of a nucleus is shown (left panel). The number of PLA dots per nucleus in cells treated with both antibodies (CTL) or with only PA28γ antibodies (w/o anti-HP1α) is shown on the bar graph (right panel). Data represent the mean ± SD from 3 independent experiments, the number of cells analyzed was *n* = 40 and *n* = 41 in control cells and cells treated without primary HP1α antibody, respectively. The *p*-value was determined using Student’s *t*-test, **** *p* ≤ 0.0001).

**Figure S5. PA28γ loss has neither a global effect on H3K9me3, H4K20me3 and H3K4me3 protein level nor on HP1β -binding and H3K4me3 mark at repetitive DNA sequences and genes.**

**A.** Representative immunoblots of whole-cell extracts from U2OS (WT and KO-PA28γ) cells, using anti-H3K9me3 antibodies. Histone H3 was used as loading control. Graphical representation of the relative abundance of the tri-methylation (H3K9me3) mark on histone H3 normalized to histone H3. The mean ± SD is from four independent experiments. The *p*-value was determined using a Student’s *t*-test, ns = not significant (*p* = 0.9354).

**B.** Immunoblots of whole-cell extracts from U2OS (WT and KO-PA28γ) cells, using anti-H4K20me3 and anti-H4K20me1 antibodies. Histone H3 was used as loading control. Graphical representation of the relative abundance of the mono-methylation (H4K20me1) and the tri-methylation (H4K20me3) marks on histone H4 normalized to histone H3. The mean ± SD is from four independent experiments. The *p*-value was determined using Student’s *t*-test, **** *p* ≤ 0.0001 *(p* = 2.091.74E-07 and *p* = 9.25E-05 for H4K20me1 andH4K20me3, respectively).

**C.** ChIP-qPCR analysis of HP1β levels at different repetitive elements and genes (as indicated on the x-axis) in WT *versus* KO-PA28γ U2OS cells. Data are represented as relative enrichment of HP1β antibody *versus* histone H3 control, as shown on the y-axis. Data are means +/− SEM (*n* = 3). Significance was calculated by Student’s *t*-test, ns = not significant, *(p* = 0.9809, *p* = 0.4746, *p* = 0.5446 and *p* = 0.5554 for HERV-K, Satll, α-Sat and GAPDH respectively), *\*p* < 0.05 *(p* = 0.01723 and *p* = 0.01763 for LINE-1 and PSMB2, respectively).

**D.** Representative immunoblots of whole-cell extracts from U2OS (WT and KO-PA28γ) cells, using anti-H3K4me3 antibodies. Histone H3 was used as loading control. Graphical representation of the relative abundance of the tri-methylation (H3K4me3) mark on histone H3 normalized to histone H3. The mean ± SD is from four independent experiments. The *p*-value was determined with a Student’s *t*-test, ns = not significant *(p* = 0.9354).

**E.** ChIP-qPCR analysis of H3K4me3 levels at different repetitive elements and genes (as indicated on the x-axis) in WT *versus* KO-PA28γ U2OS cells. Data are represented as relative enrichment of H3K4me3 *versus* histone H3 control as shown on the y-axis. Data are means +/− SEM (*n* = 3). Significance was calculated by Student’s *t*-test, ns = not significant *(p* = 0.8453, *p* = 0.8116, *p* = 0.1863 and *p* = 0.4721 for, LINE-1, GAPDH and PSMB2, respectively).

**Figure S6. PA28γ depletion decreases the S phase duration.**

Immunoblot of total cell extracts from asynchronous parental (WT) and KO-PA28γ U2OS cells (AS), and cells synchronized at the G1/S phase transition by a double thymidine block (0) and released for the times indicated. Immunoblot were probed using the antibodies indicated. β-actin was used as a loading control.

## References

Alpsoy A., Dykhuizen E. C. (2018). Glioma tumor suppressor candidate region gene 1 (GLTSCR1) and its paralog GLTSCR1-like form SWI/SNF chromatin remodeling subcomplexes. J Biol Chem 293, 3892–3903.

Ayarpadikannan S., Kim H. S. (2014). The impact of transposable elements in genome evolution and genetic instability and their implications in various diseases. Genomics Inform 12, 98–104.

Baldin V., Militello M., Thomas Y., Doucet C., Fic W., Boireau S., Jariel-Encontre I., Piechaczyk M., Bertrand E., Tazi J. et al (2008). A novel role for PA28gamma-proteasome in nuclear speckle organization and SR protein trafficking. Mol Biol Cell 19, 1706–1716.

Barton L. F., Runnels H. A., Schell T. D., Cho Y., Gibbons R., Tevethia S. S., Deepe G. S., Jr., Monaco J. J. (2004). Immune defects in 28-kDa proteasome activator gamma-deficient mice. J Immunol 172, 3948–3954.

Beck D. B., Oda H., Shen S. S., Reinberg D. (2012). PR-Set7 and H4K20me1: at the crossroads of genome integrity, cell cycle, chromosome condensation, and transcription. Genes Dev 26, 325–337.

Bosch-Presegue L., Raurell-Vila H., Thackray J. K., Gonzalez J., Casal C., Kane-Goldsmith N., Vizoso M., Brown J. P., Gomez A., Ausio J. et al (2017). Mammalian HP1 Isoforms Have Specific Roles in Heterochromatin Structure and Organization. Cell Rep 21, 2048–2057.

Brustel J., Kirstein N., Izard F., Grimaud C., Prorok P., Cayrou C., Schotta G., Abdelsamie A. F., Dejardin J., Mechali M. et al (2017). Histone H4K20 tri-methylation at late-firing origins ensures timely heterochromatin replication. Embo J 36, 2726–2741.

Chen X., Barton L. F., Chi Y., Clurman B. E., Roberts J. M. (2007). Ubiquitin-independent degradation of cell-cycle inhibitors by the REGgamma proteasome. Mol Cell 26, 843–852.

Cioce M., Boulon S., Matera A. G., Lamond A. I. (2006). UV-induced fragmentation of Cajal bodies. J Cell Biol 175, 401–413.

Collins G. A., Goldberg A. L. (2017). The Logic of the 26S Proteasome. Cell 169, 792–806.

Coux O., Zieba B. A., Meiners S. (2020). The Proteasome System in Health and Disease. Adv Exp Med Biol 1233, 55–100.

Dambacher S., Hahn M., Schotta G. (2013). The compact view on heterochromatin. Cell Cycle 12, 2925–2926.

Djeghloul D., Kuranda K., Kuzniak I., Barbieri D., Naguibneva I., Choisy C., Bories J. C., Dosquet C., Pla M., Vanneaux V. et al (2016). Age-Associated Decrease of the Histone Methyltransferase SUV39H1 in HSC Perturbs Heterochromatin and B Lymphoid Differentiation. Stem Cell Reports 6, 970–984.

Erdel F., Rippe K. (2018). Formation of Chromatin Subcompartments by Phase Separation. Biophys J 114, 2262–2270.

Erdel F., Rademacher A., Vlijm R., Tünnermann J., Frank L., Weinmann R., Schweigert E., Yserentant K., Hummert J., Bauer C. et al (2020). Mouse Heterochromatin Adopts Digital Compaction States without Showing Hallmarks of HP1-Driven Liquid-Liquid Phase Separation. Mol Cell 78, 236–249.

Fabre B., Lambour T., Garrigues L., Ducoux-Petit M., Amalric F., Monsarrat B., Burlet-Schiltz O., Bousquet-Dubouch M. P. (2014). Label-free quantitative proteomics reveals the dynamics of proteasome complexes composition and stoichiometry in a wide range of human cell lines. J Proteome Res 13, 3027–3037.

Filipponi D., Muller J., Emelyanov A., Bulavin D. V. (2013). Wip1 controls global heterochromatin silencing via ATM/BRCA1-dependent DNA methylation. Cancer Cell 24, 528–541.

Förster A., Masters E.I., Whitby F.G., Robinson H., Hill C.P. (2005). The 1.9 Å structure of a proteasome-11S activator complex and implications for proteasome-PAN/PA700 interactions. Molecular Cell 18, 589–599.

Geng F., Tansey W. P. (2012). Similar temporal and spatial recruitment of native 19S and 20S proteasome subunits to transcriptionally active chromatin. Proc Natl Acad Sci U S A 109, 6060–6065.

Grewal S. I., Jia S. (2007). Heterochromatin revisited. Nat Rev Genet 8, 35–46.

Guillot P. V., Xie S. Q., Hollinshead M., Pombo A. (2004). Fixation-induced redistribution of hyperphosphorylated RNA polymerase II in the nucleus of human cells. Exp Cell Res 295, 460–468.

Howe F. S., Fischl H., Murray S. C., Mellor J. (2017). Is H3K4me3 instructive for transcription activation?. BioEssays 39, e201600095.

Janssen A., Colmenares S. U., Karpen G. H. (2018). Heterochromatin: Guardian of the Genome. Annu Rev Cell Dev Biol 34, 265–288.

Jegou T., Chung I., Heuvelman G., Wachsmuth M., Gorisch S. M., Greulich-Bode K. M., Boukamp P., Lichter P., Rippe K. (2009). Dynamics of telomeres and promyelocytic leukemia nuclear bodies in a telomerase-negative human cell line. Mol Biol Cell 20, 2070–2082.

Jonik-Nowak B., Menneteau T., Fesquet D., Baldin V., Bonne-Andrea C., Mechali F., Fabre B., Boisguerin P., de Rossi S., Henriquet C. et al (2018). PIP30/FAM192A is a novel regulator of the nuclear proteasome activator PA28gamma. Proc Natl Acad Sci U S A 115, E6477–E6486.

Kito Y., Matsumoto M., Hatano A., Takami T., Oshikawa K., Matsumoto A., Nakayama K. I. (2020). Cell cycle-dependent localization of the proteasome to chromatin. Sci Rep 10, 5801.

Klement K., Goodarzi A. A. (2014). DNA double strand break responses and chromatin alterations within the aging cell. Exp Cell Res 329, 42–52.

Kowarz E., Loscher D., Marschalek R. (2015). Optimized Sleeping Beauty transposons rapidly generate stable transgenic cell lines. Biotechnol J 10, 647–653.

Kumar A., Kono H. (2020). Heterochromatin protein 1 (HP1): interactions with itself and chromatin components Biophysical Reviews 12, 387–400.

Lachner M., O’Carroll D., Rea S., Mechtler K., Jenuwein T. (2001). Methylation of histone H3 lysine 9 creates a binding site for HP1 proteins. Nature 410, 116–120.

Larson A. G., Elnatan D., Keenen M. M., Trnka M. J., Johnston J. B., Burlingame A. L., Agard D. A., Redding S., Narlikar G. J. (2017). Liquid droplet formation by HP1alpha suggests a role for phase separation in heterochromatin. Nature 547, 236–240.

Levy-Barda A., Lerenthal Y., Davis A. J., Chung Y. M., Essers J., Shao Z., van Vliet N., Chen D. J., Hu M. C., Kanaar R. et al (2011). Involvement of the nuclear proteasome activator PA28gamma in the cellular response to DNA double-strand breaks. Cell Cycle 10, 4300–4310.

Li S., Jiang C., Pan J., Wang X., Jin J., Zhao L., Pan W., Liao G., Cai X., Li X. et al (2015). Regulation of c-Myc protein stability by proteasome activator REGgamma. Cell Death Differ 22, 1000–1011.

Li X., Amazit L., Long W., Lonard D. M., Monaco J. J., O’Malley B. W. (2007). Ubiquitin-and ATP-independent proteolytic turnover of p21 by the REGgamma-proteasome pathway. Mol Cell 26, 831–842.

Li X., Lonard D. M., Jung S. Y., Malovannaya A., Feng Q., Qin J., Tsai S. Y., Tsai M. J., O’Malley B. W. (2006). The SRC-3/AIB1 coactivator is degraded in a ubiquitin-and ATP-independent manner by the REGgamma proteasome. Cell 124, 381–392.

Lippman Z., Gendrel A. V., Black M., Vaughn M. W., Dedhia N., McCombie W. R., Lavine K., Mittal V., May B., Kasschau K. D. et al (2004). Role of transposable elements in heterochromatin and epigenetic control. Nature 430, 471–476.

Liu Y., Qin S., Lei M., Tempel W., Zhang Y., Loppnau P., Li Y., Min J. (2017). Peptide recognition by heterochromatin protein 1 (HP1) chromoshadow domains revisited: Plasticity in the pseudosymmetric histone binding site of human HP1 J Biol Chem 292, 5655–5664.

Lleres D., James J., Swift S., Norman D. G., Lamond A. I. (2009). Quantitative analysis of chromatin compaction in living cells using FLIM-FRET. J Cell Biol 187, 481–496.

Ma C. P., Slaughter C. A., DeMartino G. N. (1992). Identification, purification, and characterization of a protein activator (PA28) of the 20 S proteasome (macropain). J Biol Chem 267, 10515–10523.

Ma C. P., Willy P. J., Slaughter C. A., DeMartino G. N. (1993). PA28, an activator of the 20 S proteasome, is inactivated by proteolytic modification at its carboxyl terminus. J Biol Chem 268, 22514–22519.

Machida S., Takizawa Y., Ishimaru M., Sugita Y., Sekine S., Nakayama J. I., Wolf M., Kurumizaka H. (2018). Structural Basis of Heterochromatin Formation by Human HP1. Mol Cell 69, 385–397 e388.

Maison C., Almouzni G. (2004). HP1 and the dynamics of heterochromatin maintenance. Nat Rev Mol Cell Biol 5, 296–304.

Mao I., Liu J., Li X., Luo H. (2008). REGgamma, a proteasome activator and beyond? Cell Mol Life Sci 65, 3971–3980.

Martin C., Zhang Y. (2005). The diverse functions of histone lysine methylation. Nat Rev Mol Cell Biol 6, 838–849.

Masson P., Lundgren J., Young P. (2003). Drosophila proteasome regulator REGgamma: transcriptional activation by DNA replication-related factor DREF and evidence for a role in cell cycle progression. J Mol Biol 327, 1001–1012.

McCann T. S., Tansey W. P. (2014). Functions of the proteasome on chromatin. Biomolecules 4, 1026–1044.

Murata S., Kawahara H., Tohma S., Yamamoto K., Kasahara M., Nabeshima Y., Tanaka K., Chiba T. (1999). Growth retardation in mice lacking the proteasome activator PA28gamma. J Biol Chem 274, 38211–38215.

Nishibuchi G., Nakayama J. (2014). Biochemical and structural properties of heterochromatin protein 1: understanding its role in chromatin assembly. J Biochem 156, 11–20.

Oda H., Okamoto I., Murphy N., Chu J., Price S. M., Shen M. M., Torres-Padilla M. E., Heard E., Reinberg D. (2009). Monomethylation of histone H4-lysine 20 is involved in chromosome structure and stability and is essential for mouse development. Mol Cell Biol 29, 2278–2295.

Otterstrom J., Castells-Garcia A., Vicario C., Gomez-Garcia P. A., Cosma M. P., Lakadamyali M. (2019). Super-resolution microscopy reveals how histone tail acetylation affects DNA compaction within nucleosomes in vivo. Nucleic Acids Res 47, 8470–8484.

Padeken J., Zeller P., Gasser S. M. (2015). Repeat DNA in genome organization and stability. Curr Opin Genet Dev 31, 12–19.

Peng J. C., Karpen G. H. (2009). Heterochromatic genome stability requires regulators of histone H3 K9 methylation PLoS Genet 5, e1000435.

Peters A. H., O’Carroll D., Scherthan H., Mechtler K., Sauer K. S., Schöfer C., Weipoltshammer K., Pagani M., M Lachner M., Kohlmaier A. et al (2001). Loss of the Suv39h histone methyltransferases impairs mammalian heterochromatin and genome stability Cell 107, 323–327.

Price B. D., D’Andrea A. D. (2013). Chromatin remodeling at DNA double-strand breaks. Cell 152, 1344–1354.

Rechsteiner M., Hill C. P. (2005). Mobilizing the proteolytic machine: cell biological roles of proteasome activators and inhibitors. Trends Cell Biol 15, 27–33.

Saksouk N., Hajdari S., Perez Y., Pratlong M., Barrachina C., Graber C., Grégoire D., Zavoriti A., Sarrazi A., Pirot N. et al (2020). The mouse HP1 proteins are essential for preventing liver tumorigenesis Oncogene 39, 2676–2691.

Saksouk N., Simboeck E., Dejardin J. (2015). Constitutive heterochromatin formation and transcription in mammals. Epigenetics Chromatin 8, 3.

Sawyer I. A., Bartek J., Dundr M. (2019). Phase separated microenvironments inside the cell nucleus are linked to disease and regulate epigenetic state, transcription and RNA processing. Semin Cell Dev Biol 90, 94–103.

Schotta G., Lachner M., Sarma K., Ebert A., Sengupta R., Reuter G., Reinberg D., Jenuwein T. (2004). A silencing pathway to induce H3-K9 and H4-K20 trimethylation at constitutive heterochromatin. Genes Dev 18, 1251–1262.

Shoaib M., Walter D., Gillespie P. J., Izard F., Fahrenkrog B., Lleres D., Lerdrup M., Johansen J. V., Hansen K., Julien E. et al (2018). Histone H4K20 methylation mediated chromatin compaction threshold ensures genome integrity by limiting DNA replication licensing. Nat Commun 9, 3704.

Soderberg O., Gullberg M., Jarvius M., Ridderstrale K., Leuchowius K. J., Jarvius J., Wester K., Hydbring P., Bahram F., Larsson L. G. et al (2006). Direct observation of individual endogenous protein complexes in situ by proximity ligation. Nat Methods 3, 995–1000.

Strom A. R., Emelyanov A. V., Mir M., Fyodorov D. V., Darzacq X., Karpen G. H. (2017). Phase separation drives heterochromatin domain formation. Nature 547, 241–245.

Sun L., Fan G., Shan P., Qiu X., Dong S., Liao L., Yu C., Wang T., Gu X., Li Q. et al (2016). Regulation of energy homeostasis by the ubiquitin-independent REGgamma proteasome. Nat Commun 7, 12497.

Tardat M., Murr R., Herceg Z., Sardet C., Julien E. (2007). PR-Set7-dependent lysine methylation ensures genome replication and stability through S phase. J Cell Biol 179, 1413–1426.

Thiru A., Nietlispach D., Mott H. R., Okuwaki M., Lyon D., Nielsen P. R., Hirshberg M., Verreault A., Murzina N. V., Laue E. D. (2004). Structural basis of HP1/PXVXL motif peptide interactions and HP1 localisation to heterochromatin. Embo J 23, 489–499.

Thomas Y., Peter M., Mechali F., Blanchard J. M., Coux O., Baldin V. (2014). Kizuna is a novel mitotic substrate for CDC25B phosphatase. Cell Cycle 13, 3867–3877.

Verschure P. J., van der Kraan I., de Leeuw W., van der Vlag J., Carpenter A. E., Belmont A. S., van Driel R. (2005). In vivo HP1 targeting causes large-scale chromatin condensation and enhanced histone lysine methylation. Mol Cell Biol 25, 4552–4564.

Welk V., Coux O., Kleene V., Abeza C., Trumbach D., Eickelberg O., Meiners S. (2016). Inhibition of Proteasome Activity Induces Formation of Alternative Proteasome Complexes. J Biol Chem 291, 13147–13159.

Wilk S., Chen W. E., Magnusson R. P. (2000). Properties of the nuclear proteasome activator PA28gamma (REGgamma). Arch Biochem Biophys 383, 265–271.

Wojcik C., Tanaka K., Paweletz N., Naab U., Wilk S. (1998). Proteasome activator (PA28) subunits, alpha, beta and gamma (Ki antigen) in NT2 neuronal precursor cells and HeLa S3 cells. Eur J Cell Biol 77, 151–160.

Zannini L., Buscemi G., Fontanella E., Lisanti S., Delia D. (2009). REGgamma/PA28gamma proteasome activator interacts with PML and Chk2 and affects PML nuclear bodies number. Cell Cycle 8, 2399–2407.

Zannini L., Lecis D., Buscemi G., Carlessi L., Gasparini P., Fontanella E., Lisanti S., Barton L., Delia D. (2008). REGgamma proteasome activator is involved in the maintenance of chromosomal stability. Cell Cycle 7, 504–512.

Zeng W., Ball A. R., Jr., Yokomori K. (2010). HP1: heterochromatin binding proteins working the genome. Epigenetics 5, 287–292.

Zhang Z., Zhang R. (2008). Proteasome activator PA28 gamma regulates p53 by enhancing its MDM2-mediated degradation. Embo J 27, 852–864.

