## Supplementary Figures for "The 20S proteasome activator PA28γ controls the compaction of chromatin"

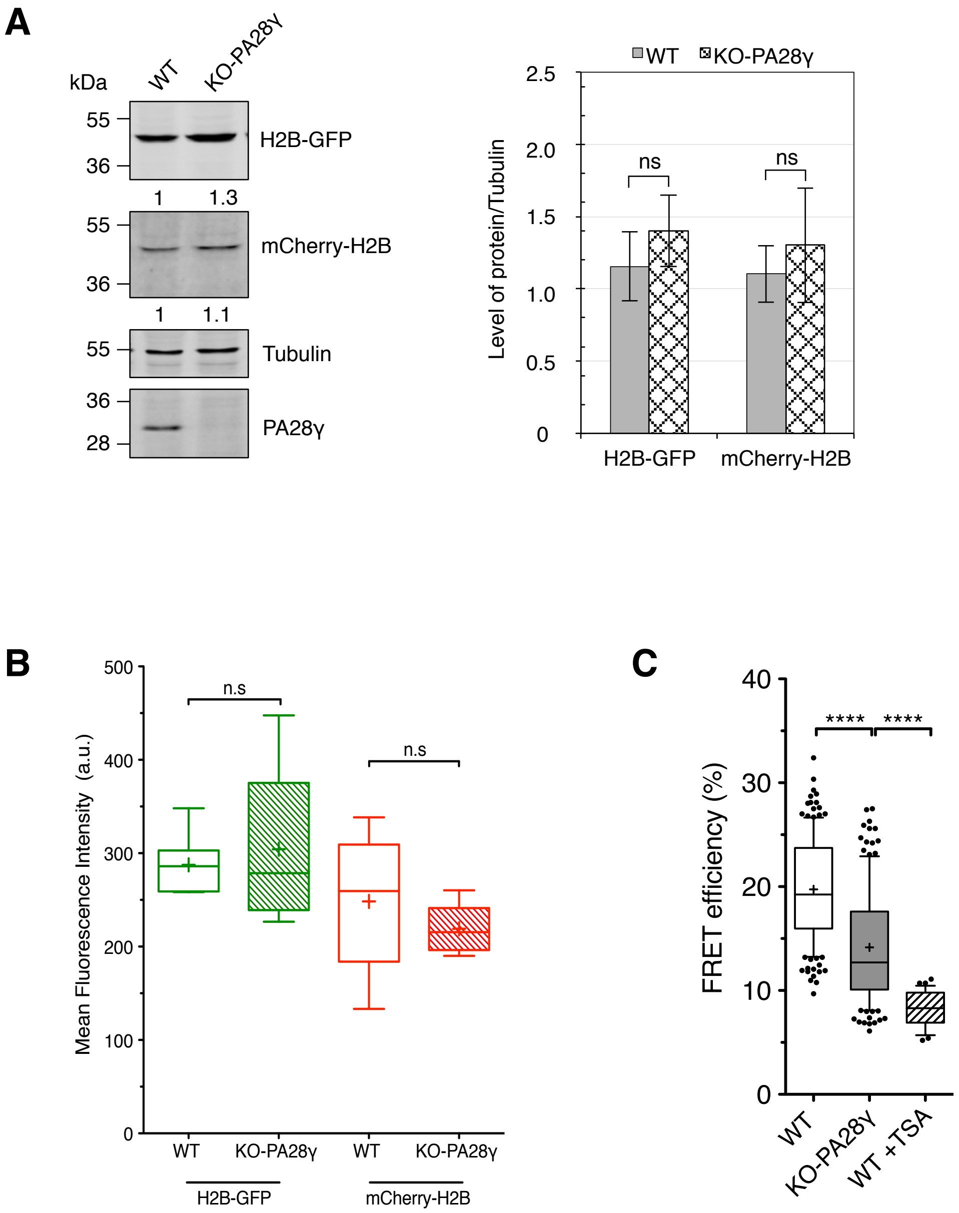
A
kDa
WT
KO-PA28γ
55
36
H2B-GFP
1
1.3
55
36
mCherry-H2B
1
1.1
55
Tubulin
36
28
PA28γ
WT
KO-PA28γ
2.5
2.0
1.5
1.0
0.5
0
ns
ns
Level of protein/Tubulin
H2B-GFP
mCherry-H2B
B
500
400
300
200
100
0
n.s
n.s
Mean Fluorescence Intensity (a.u.)
WT
KO-PA28γ
WT
KO-PA28γ
H2B-GFP
mCherry-H2B
C
40
30
20
10
0
****
****
FRET efficiency (%)
WT
KO-PA28γ
WT +TSA

Figure S1

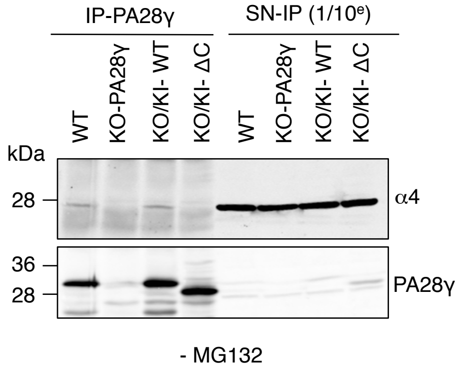
IP-PA28γ
SN-IP (1/10e)
WT
KO-PA28γ
KO/KI- WT
KO/KI- ΔC
WT
KO-PA28γ
KO/KI- WT
KO/KI- ΔC
kDa
28
α4
36
28
PA28γ
- MG132

Figure S2

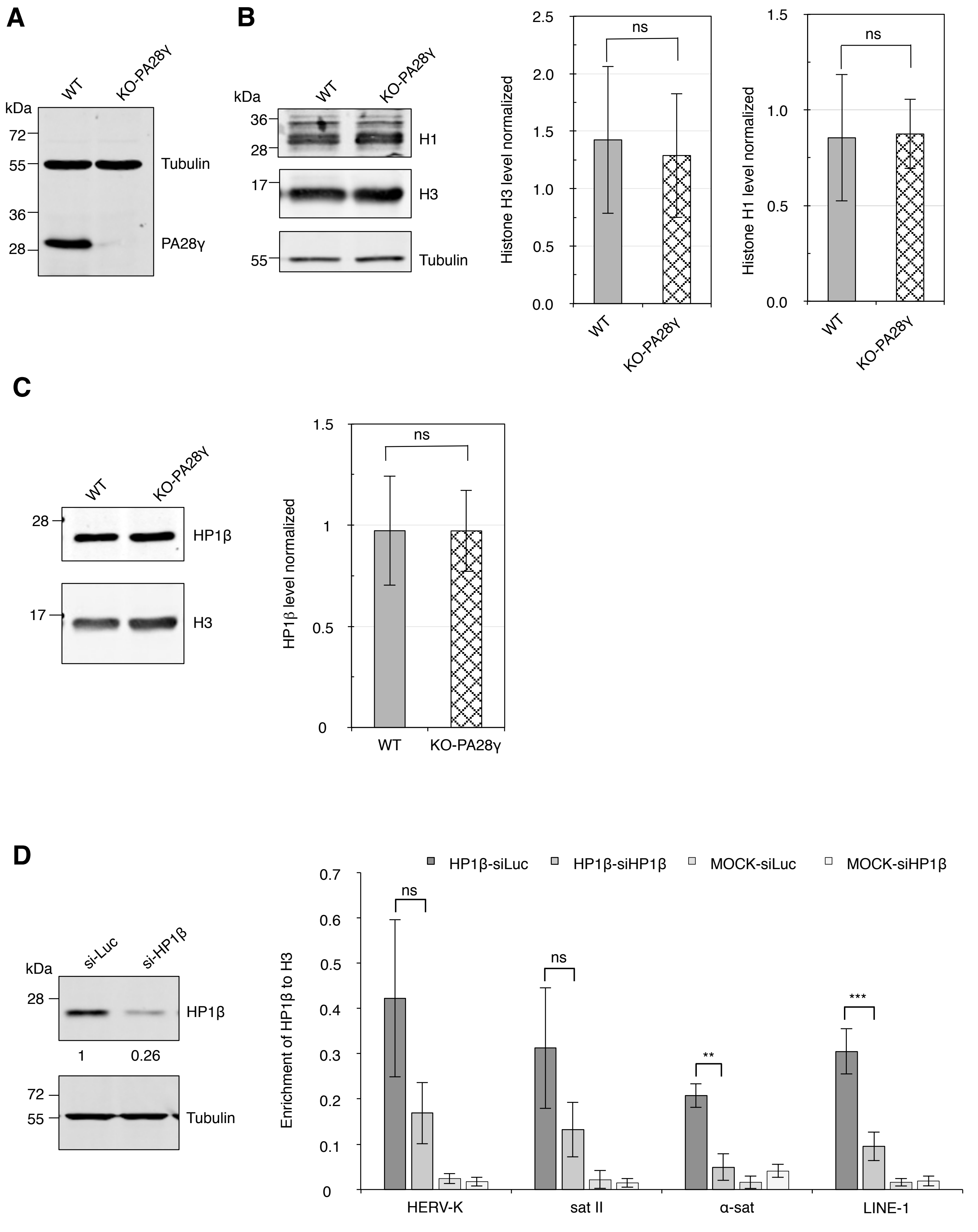
A
WT
KO-PA28γ
kDa
72
55
36
28
Tubulin
PA28γ
B
WT
KO-PA28γ
kDa
36
28
17
55
H1
H3
Tubulin
Histone H3 level normalized
2.5
2.0
1.5
1.0
0.5
0.0
ns
WT
KO-PA28γ
Histone H1 level normalized
1.5
1.0
0.5
0.0
ns
WT
KO-PA28γ
C
WT
KO-PA28γ
28
17
HP1β
H3
HP1β level normalized
1.5
1
0.5
0
ns
WT
KO-PA28γ
D
si-Luc
si-HP1β
kDa
28
HP1β
1
0.26
72
55
Tubulin
HP1β-siLuc
HP1β-siHP1β
MOCK-siLuc
MOCK-siHP1β
Enrichment of HP1β to H3
0.7
0.6
0.5
0.4
0.3
0.2
0.1
0
ns
ns
**
***
HERV-K
sat II
α-sat
LINE-1

Figure S3

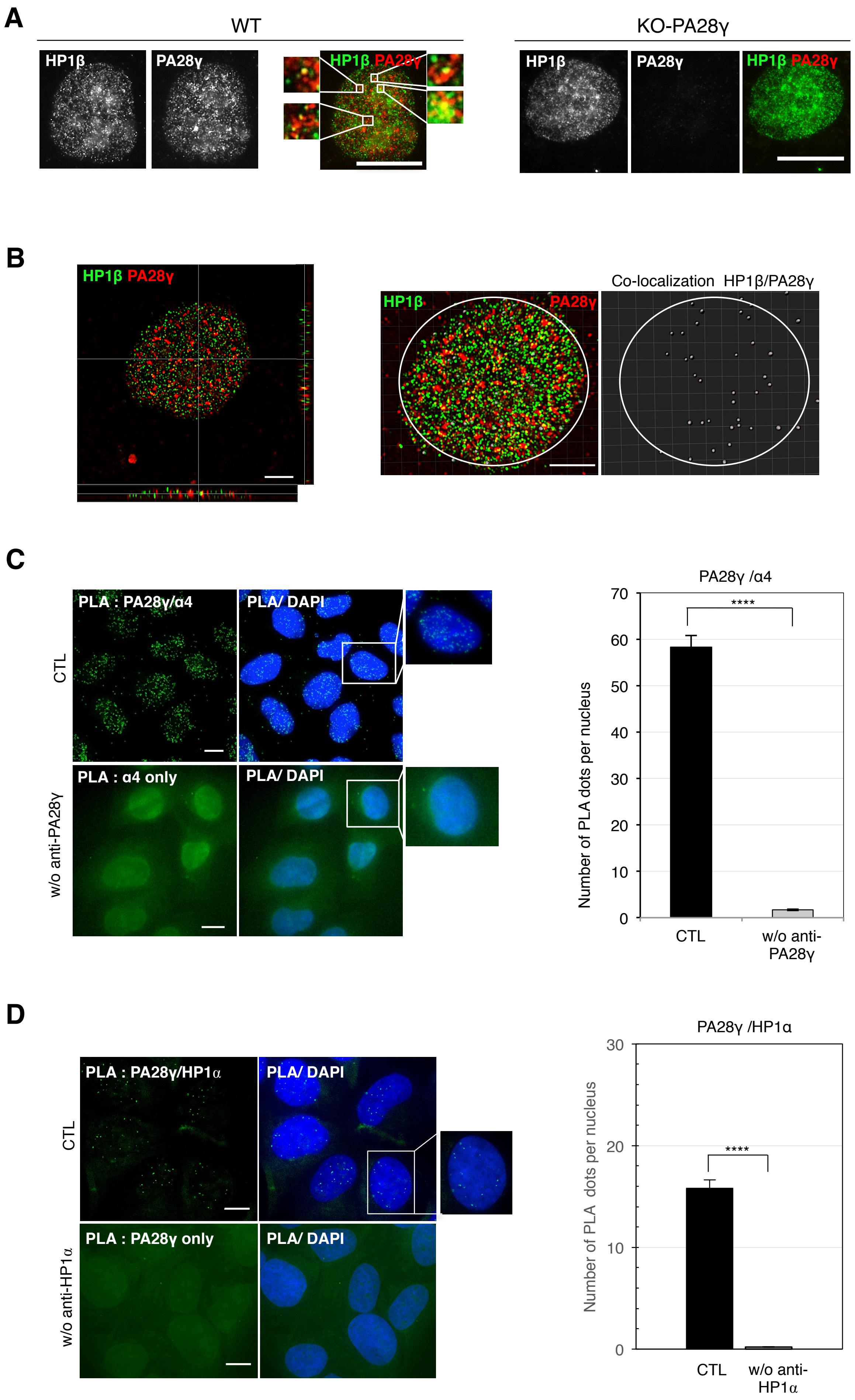
A
WT
KO-PA28γ
HP1β
PA28γ
HP1β PA28γ
B
HP1β PA28γ
Co-localization HP1β/PA28γ
C
PLA : PA28γ/α4
PLA/ DAPI
CTL
PLA : α4 only
w/o anti-PA28γ
PA28γ /α4
Number of PLA dots per nucleus
****
CTL
w/o anti-PA28γ
D
PLA : PA28γ/HP1α
PLA/ DAPI
CTL
PLA : PA28γ only
w/o anti-HP1α
PA28γ /HP1α
Number of PLA dots per nucleus
****
CTL
w/o anti-HP1α

Figure S4

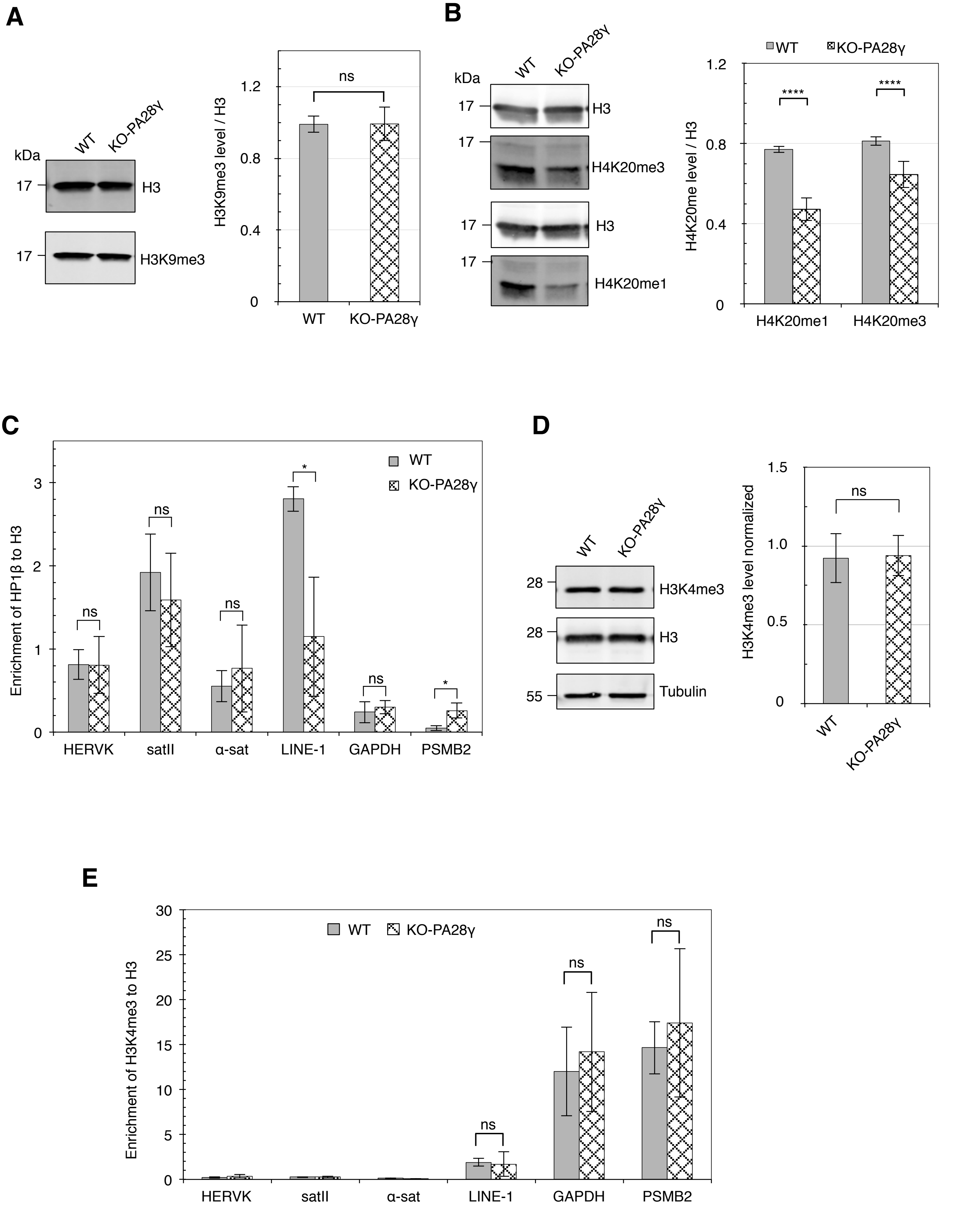
A
kDa
WT
KO-PA28γ
17
H3
17
H3K9me3
H3K9me3 level / H3
ns
1.2
0.8
0.4
0
WT
KO-PA28γ
B
kDa
WT
KO-PA28γ
17
H3
17
H4K20me3
17
H3
17
H4K20me1
WT
KO-PA28γ
H4K20me level / H3
****
****
1.2
0.8
0.4
0
H4K20me1
H4K20me3
C
Enrichment of HP1β to H3
WT
KO-PA28γ
ns
ns
ns
*
ns
*
3
2
1
0
HERVK
satII
α-sat
LINE-1
GAPDH
PSMB2
D
WT
KO-PA28γ
28
H3K4me3
28
H3
55
Tubulin
H3K4me3 level normalized
ns
1.5
1.0
0.5
0
WT
KO-PA28γ
E
Enrichment of H3K4me3 to H3
WT
KO-PA28γ
ns
ns
ns
30
25
20
15
10
5
0
HERVK
satII
α-sat
LINE-1
GAPDH
PSMB2

Figure S5

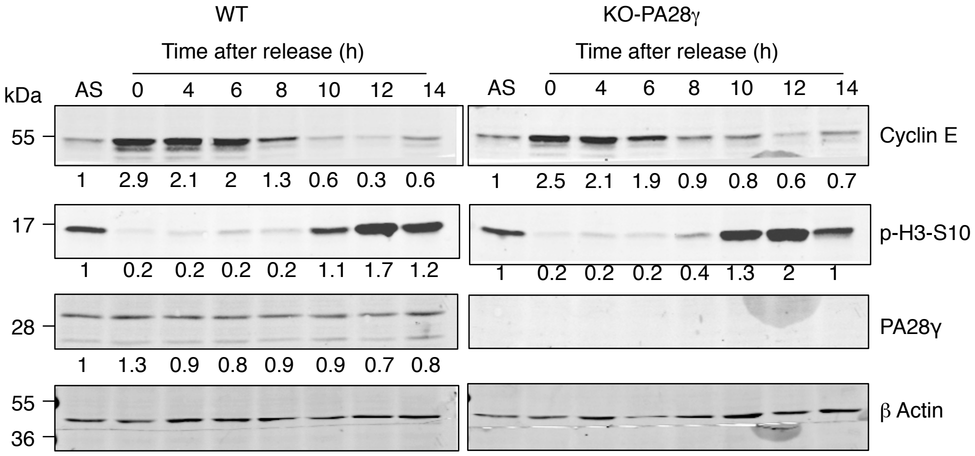
WT
KO-PA28γ
Time after release (h)
Time after release (h)
kDa
AS 0 4 6 8 10 12 14
AS 0 4 6 8 10 12 14
55
Cyclin E
1 2.9 2.1 2 1.3 0.6 0.3 0.6
1 2.5 2.1 1.9 0.9 0.8 0.6 0.7
17
p-H3-S10
1 0.2 0.2 0.2 0.2 1.1 1.7 1.2
1 0.2 0.2 0.2 0.4 1.3 2 1
28
PA28γ
1 1.3 0.9 0.8 0.9 0.9 0.7 0.8
55
36
β Actin

Figure S6

**Table S1**: *p* values of the figure 6A,B (ChIP)

| | T-test (H3K9me3/H3) U2OS *vs* KO PA28γ | T-test (H4K20me3/H3) U2OS *vs* KO PA28γ |
|---|---|---|
| HERV-K | 6.47871E-06 | 0.009044059 |
| Sat II | 8.6538E-06 | 1.37015E-05 |
| α-sat | 0.00014585 | 0.001004765 |
| LINE-1 | 2.80855E-05 | 0.002330735 |
| GAPDH | 0.677649943 | 0.003366684 |
| PSMB2 | 0.000677637 | 0.121036886 |
| CCNA2 | 0.848275718 | 0.066976205 |
| CCNE2 | 0.803674003 | 0.835817889 |

**Table S2**: *p* values of the figure S3D (ChIP)

| | T-test (HP1β) U2OS si-Luc *vs* si-HP1β) |
|---|---|
| HERV-K | 0.057078569 |
| Sat II | 0.066693611 |
| α-sat | 0.005250998 |
| LINE-1 | 0.000382203 |

**Table S3**: *p* values of the figure 6C (determined with the 2-way ANOVA)

| Time after release (hours) | 0 | 4 | 6 | 8 | 10 | 12 |
|---|---|---|---|---|---|---|
| S-phase | 0.6759 | 0.0052 | < 0.0001 | < 0.0001 | > 0.9999 | 0.0377 |
| G2/M-phase | 0.9946 | 0.0187 | < 0.0001 | < 0.0001 | < 0.0002 | < 0.0001 |
